# The mutational dynamics of the Arabidopsis centromeres

**DOI:** 10.1101/2025.06.02.657473

**Authors:** Xiao Dong, Wen-Biao Jiao, Lara Goldkuhle, Fernando Rabanal, Samija Amar, Matthew T. Parker, José A Campoy, Yueqi Tao, Bruno Huettel, Jurriaan Ton, Lisa M. Smith, Holger Puchta, Detlef Weigel, Korbinian Schneeberger

## Abstract

Centromeres are specialized chromosomal regions essential for sister chromatid cohesion and spindle attachment during cell division. Although many centromeres consist of highly variable tandem repeat arrays, the mutational processes driving this variability remain poorly understood. Here, using replicated genome assemblies of *Arabidopsis thaliana* mutation accumulation (MA) lines, we define the centromeric mutation spectrum. We find that point mutations occur at an almost tenfold higher rates than in chromosome arms, largely driven by non-allelic gene conversion between closely linked repeat units. Large kilobase-scale indels are also frequent and consistently preserve the tandem repeat array by adding or removing only complete repeat units. Analysis of MA lines deficient in the helicase RTEL1 supports the involvement of homology-directed DNA repair in these mutational processes. Moreover, simulations of sequence turnover using the centromere-specific mutation spectrum recapitulate the emergence of homogenized repeat blocks characteristic of natural centromeres. Together, our results show that centromere evolution is driven by a distinct mutational spectrum shaped by homology-directed DNA repair, providing a quantitative framework for how local mutational processes generate and maintain the large-scale architecture of centromeric DNA.

## Introduction

Centromeres are essential for accurate chromosome segregation during cell division. Despite its conserved function, centromeric DNA often consists of megabase-scale satellite repeat arrays that vary markedly both within and between species^1–5^. However, the mutational processes driving this rapid sequence turnover remain poorly understood.

The individual repeat units of centromeric satellite repeat arrays are usually quite small (∼100 to 200 base pairs)^6,7^. They are similar in sequence, but not identical to each other. The sequence differences between them generate distinct patterns in the satellite array, such as tandem duplications or triplications. These patterns can be further extended into higher-order repeat (HORs) structures, where the same repeat unit patterns are repeated many times. If they are similar, closely-linked HORs form homogenized blocks that can span several megabases in size and that underly the characteristic patterns commonly observed with centromere repeat visualisation^1,5,6^. The functional cores of the centromeric repeat arrays often consist of such homogenized blocks, flanked by heterogeneous peripheries, which are characterized by structural rearrangements and transposable element insertions^1,5^. While this general structure is conserved between different accessions within a species, the actual arrangement of the homogenized blocks, their number and sizes are highly variable, which has led to the hypothesis that megabase-scale mutations or long-range recombination events are required to form centromeric sequences^7^.

In *A. thaliana*, the five centromeres consist of megabase-scale tandem arrays of derivatives of a 178 bp repeat monomer (*CEN178*)^5^. Long-read assemblies have successfully reconstructed these arrays across all five chromosomes within the reference line and across large populations^5,8–10^. The extreme sequence divergence observed in natural centromeres supports the hypothesis that these regions are shaped by fundamentally different, so-far unknown mutational dynamics when compared to the rest of the genome^1,11^.

To better understand these mutational processes, we analysed *A. thaliana* mutation accumulation (MA) lines, which are propagated through repeated single-seed descent for several generations to allow mutations to accumulate. However, even though centromeres can now be assembled^5,8–10^, identifying rare mutations in the centromeres of MA lines remains challenging due to the highly repetitive nature of the centromeric DNA that impacts the quality of the assemblies. To overcome this, we developed an innovative replicated genome sequencing strategy to eliminate genome assembly errors, allowing us to distinguish genuine centromeric mutations from assembly artefacts.

Applying this approach to several MA lines revealed a unique mutational spectrum in the centromeres. The mutations included frequent tandem-repeat-preserving indels of several kb in size that added or deleted only complete repeat units, while keeping the repeat array structure intact. In addition, we found a close to tenfold enrichment of point mutations, including some clustered point mutations that were likely introduced by non-allelic gene conversions. Using MA lines derived from a mutant of the DNA helicase RTEL1, we identified *RTEL1* being implicated in the stability of plant centromeric repeat arrays and highlight the role of homology-directed repair as a key driver of centromere evolution. Even when including accessions from a “natural MA experiment” that accumulated mutations for ∼400 years, we could not find transposable elements activity in the centromeres. In-line with this, forward-in-time simulations demonstrated that the kilobase-scale indels and non-allelic gene conversions are sufficient to megabase-scale homogenized blocks, but require rare megabase-scale deletions to balance centromere size. Taken together, our findings uncover mutational dynamics that could explain how centromeres evolve their distinctive structures and how those diverge over time.

## Results

### Replication resolves assembly errors

Despite significant progress in recent years, *de novo* genome assemblies still contain errors, making it difficult to confidently identify mutations in highly repetitive regions. To find such mutations, we generated replicated genome assemblies (*i.e.*, independent assemblies from different DNA samples of the same individuals) to first remove mistakes in the assemblies, and then used these virtually error-free assemblies to distinguish true mutations from assembly errors.

As a proof of concept, we generated three independent genome assemblies each for two *A. thaliana* Col-0 plants using DNA extracted from pooled progeny (Fig. 1a). Both plants were from the progeny of the same Col-0 founder plant but had been separately propagated by self-pollination and single-seed descent for 16 generations as part of a controlled trans-generational mutation accumulation experiment. We refer to the two plants as MA16 individuals, and to the six assemblies as “A1”, “A2, “A3”, “B1”, “B2” and “B3” to distinguish both the identity of the plants (“A” or “B” plant) and the replication of the whole-genome assembly that has been performed independently three times for each of the two plants (“1”, “2” or “3”) (Fig. 1a). In general, Col-0 is a diploid, inbred and selfing plant with a homozygous genome, which is identical between different Col-0 plants, except for the mutations that occurred after individual lineages were split.

**Fig. 1.**
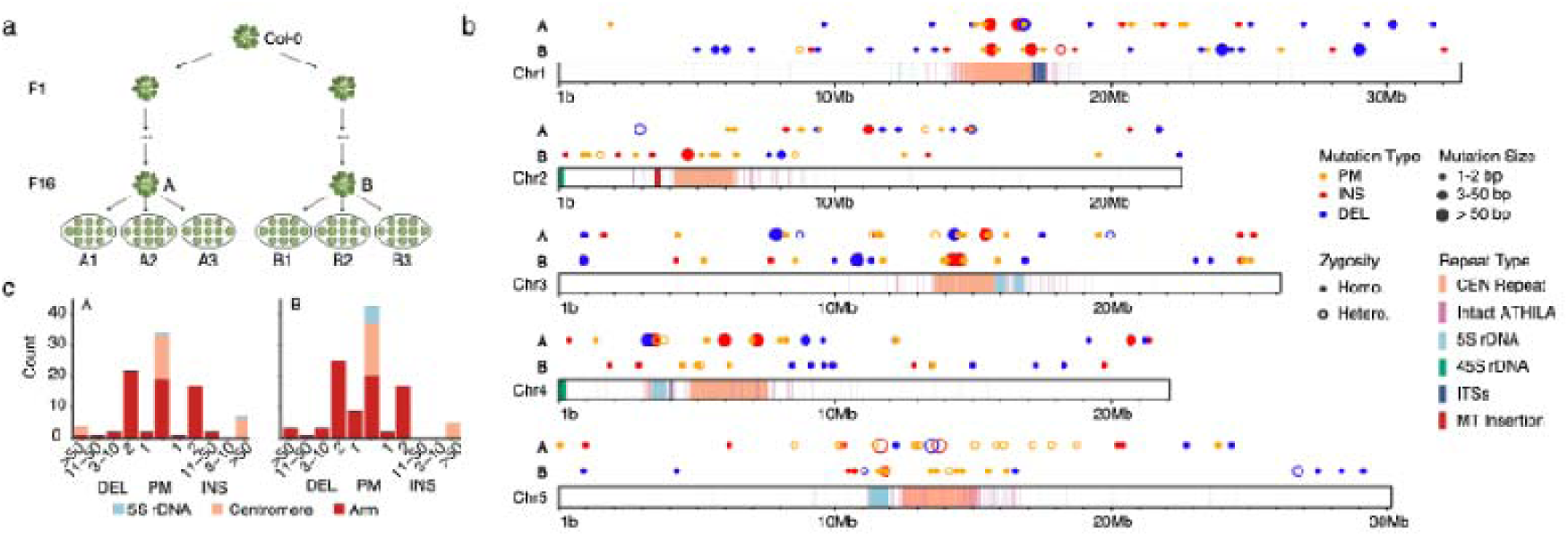
Mutation accumulation in Arabidopsis revealed by virtually error-free genome assemblies. **a.** Pools of sister plants (∼15 individuals per pool) were sequenced, with two independent pools generated from the progeny of each mother plant (designated A or B). Pooling sister genomes dilutes somatic mutations and reconstructs the genotype of the maternal plant (F16). Sequence differences between assemblies of sister pools therefore represents from assembly errors. **b.** Distribution of true mutations in samples A and B. Two rows of circles depict mutation positions, colour-coded by type: point mutations (orange), insertions (red), and deletions (blue). Circle size reflects mutation size: small (1-2 bp), medium (3-50 bp), and large (>50 bp). Solid circles mark homozygous mutations; open circles indicate heterozygous mutations. Chromosomes are shown as rectangles with repetitive regions highlighted: centromere (peach), intact ATHILA elements (pink), 5S rDNA (light blue), 45S rDNA (green), interstitial telomeric sequences (dark blue) and mitochondrial insertions (red). **c.** Bar plots showing mutation counts in samples A and B. The central bar denotes point mutations, bars to the right indicate insertions, and bars to the left deletions. Mutation size increases from the centre outward. Colours indicate genomic context: 5S rDNA (light blue), centromere (peach), and all remaining regions (red). PM: point mutation; INS: insertion; DEL: deletion; CEN, centromere; ITSs, interstitial telomeric sequences; MT, mitochondrial sequences.

Two of the assemblies of each plant were generated with PacBio HiFi (replicates “1” and “2”) and one with Oxford Nanopore Technologies (ONT) sequencing (replicate “3”). We sequenced each of the samples to very high coverage ranging from 96x to 142x (Supplementary Table 1). The PacBio assemblies featured NG50 values of 9.4 to 14.3 Mb and only 19 to 31 contigs per assembly, which were scaffolded into five pseudo-chromosomes using homology to the reference sequence (Supplementary Table 2). The ONT assemblies exhibited an even higher contiguity and included only a single assembly gap per genome implying that four of the five chromosomes were assembled into single contigs. While this suggests that the chromosomes were fully assembled, none of the six assemblies included the heterochromatic nucleolus organiser regions (NOR) at the beginning of chromosomes 2 and 4, which so far cannot been assembled by automated assembly approaches, and were only assembled with a manually resolved once for a single genome^12^.

To identify assembly errors, we compared the assemblies to each other and visually inspected every assembly difference using read alignments (Methods). While this does not resolve systematic errors that occur reproducibly across all replicates, we confirmed each of the mutations with read alignments. When comparing the replicated assemblies, we identified 391, 577, 266, and 298 errors in the PacBio assemblies (A1, A2, B1 and B2, respectively) and 10,974 and 10,801 errors in the ONT assemblies (in A3 and B3) (Extended Data Fig. 1).

The low error rate in the PacBio assemblies was confirmed by Merqury^13^, which estimated quality values ranging from 58 to 62. The most common source of errors (∼80% of all errors) in the PacBio assemblies were simple sequence repeats, where HiFi reads failed to accurately capture the correct number of repeat units and in consequence introduced small 1-2 bp indel errors into the assembly (Supplementary Fig. 1-3). Around 17% of the errors were associated with stochastic sequencing errors near GA(A) repeats, where low coverage hindered error correction through read consensus (Supplementary Fig. 4)^9^. The remaining ∼3% errors resulted from assembly artefacts introduced during reference-based scaffolding (Supplementary Fig. 5-7). Correcting the errors with conventional polishing approaches (based on the alignments of short and long sequencing reads) identified less than a quarter of all errors and at the same time “over-corrected” several hundred positions, leading to more additional than corrected errors (Supplementary Fig. 8). A detailed description of these error types is provided in the Supplementary Results (Supplementary Table 3-6).

The error profiles in the ONT assemblies were different. Nearly all of the errors in the ONT assemblies (>99.9%) consisted of small indels (<10 bp) predominately within poly-A and poly-T homopolymers with a strong bias towards deletion errors (Extended Data Fig. 1; Supplementary Fig. 9; Supplementary Table 7-8). In contrast to the PacBio assemblies, the ONT assemblies did not show lower coverage or assembly errors near GA(A) repeats, which explains the higher contiguity of these assemblies (Supplementary Fig. 10). To mitigate ONT-specific systematic errors, the assemblies were further polished using the ONT-specific tool Dorado, which substantially reduced the overall error burden. In contrast to the PacBio assemblies, polishing had a positive impact and reduced the error counts to 3,354 and 5,285 in each of the assemblies (Supplementary Fig. 11). Indel errors within simple sequence repeats, however, remained the most prevalent error type (Supplementary Table 9-10).

Across the six assemblies, 26 of the 30 centromeres were assembled into single contigs. Unexpectedly, the centromeric sequences were among the most accurately assembled regions in the genome: although they constitute 8.3% of the genome, they accounted for less than 1% of all assembly errors in both assembly types. In total, we detected only 15 assembly errors in the centromeres, including nine large (>50 bp) and six small (1-2 bp) errors. The large errors were primarily due to mis-scaffolding of the PacBio assembly contigs, while the small errors were associated with simple sequence repeats. This high level of accuracy is likely due to a marked depletion of simple sequence repeats within centromeres, which were the main source of assembly errors elsewhere in the genome (Supplementary results; Extended Data Fig. 1a).

Together, these results demonstrate that Arabidopsis centromeres can be assembled at extremely high accuracy using either PacBio or ONT read data. We used replicated genome assembly to identify and remove assembly errors allowing us to identify remaining assembly errors. In turn and in contrast to resequencing, genome assembly provides a robust approach for identifying centromeric mutations.

### Mutation identification with whole-genome assemblies

By aligning the error-corrected assemblies of different samples against each other, one can find mutations of any type in any part of the assembled genome. As heterozygous mutations can manifest in the assemblies as well, we needed to confirm the homozygosity of the assembled variants by using read alignments of the raw sequencing data against the assemblies. Here we focused on only homozygous mutations. Finally, we could assign the mutations to individual plants by using the wild-type alleles of the reference sequence.

Between the genomes of the two MA16 lines, we identified 200 homozygous mutations comprising 77 point mutations and 123 indels from 1 to 11,570 bp (Fig. 1b, c; Supplementary Table 11). Using the Col-0 reference genome, we assigned 92 mutations to plant A and 108 mutations to plant B.

To verify the completeness of the mutation calls, we compared them to earlier mutation rate estimates in Arabidopsis. Previous studies have estimated a point mutation rate of 7.0 × 10^-9^ ± 2.8 × 10^−9^ per site and generation^14,15^. The 77 homozygous point mutations (34 in A and 43 in B) are more than twice as many as expected given this mutation rate (3.9 and 5.6 standard deviations above mean). It is important to note, however, that the original estimate was based on “unique” regions of the genome (*i.e.,* regions accessible with short-read alignments) and did not include repetitive regions of the genome. In fact, when excluding the centromeres and 5S rDNA clusters (i.e., 119.8 Mb of unique regions), the expected number of mutations based on the published mutation rates was only 13.3 ± 5.3. The observed numbers of chromosome arm mutations (19 in A and 20 in B) fell well within the estimated distribution (1.07 and 1.26 standard deviations above mean), suggesting that mutation rates in the chromosome arms are consistent with previous estimates. In the chromosome arms, we can further examine transposable elements (TEs), where previous work had reported a higher mutation rate (1.4 × 10^−8^)^14,15^. Given the size of all TEs in the chromosome arms (∼4.3 Mb), we expected 1.3 and 1.4 mutations, respectively, in TEs in the chromosome arms for the two MA16 lines, which is in line with our observations of 1 mutation in A and 2 mutations in B, and further supports the accuracy of our mutation detection.

The most common type of mutation in the unique regions of the genome, however, were 2-bp indels. These small indels were exclusively found in dinucleotide repeats within the chromosome arms, which is again consistent with previous reports showing that mutation rates in dinucleotide repeats are orders of magnitude higher than those of point mutations^15,16^. All other small indels (1-50 bp) also occurred within simple sequence repeats, including three cases with slightly larger repeat units of up to 25 bp.

The remaining 38 homozygous point mutations (49%) were located within just 13.8 Mb (9.8%) of the genome, corresponding to the highly repetitive 5S rDNA clusters and the centromeres. This suggested an elevated point mutation rate in these regions compared to unique regions and indicates that these regions undergo different mutational dynamics as previously described for pericentromeric regions^15^, adding to the growing evidence that mutation rates are not uniformly distributed across the genome^15,17^.

Likewise, the majority of the large indels (>50 bp; 15 out of 19) was not associated with simple sequence repeats but occurred in highly repetitive regions such as the centromeres or the 5S rDNA clusters (Fig. 1c). Only four large deletions occurred outside these regions; with three sharing significant sequence similarity with closely linked regions, implying that local similarity is a major driver of large indel formation across the genome.

Taken together, with whole-genome assemblies, we can confidently identify mutations of any type across all assembled regions of the genome including the centromeres.

### Mutation spectrum in the centromere

To enable accurate estimation of mutation spectra and frequencies and to avoid potential biases arising from using the same genomes for both method development and mutation identification, we generated high-quality genome assemblies for eight additional MA lines that were propagated for 32 generations. As before, assemblies were produced using a replicated approach with both PacBio and ONT sequencing. In total, this dataset represents 256 generations of accumulated mutations.

Across the 40 centromeres of these eight MA32 lines, we identified 270 homozygous mutations, including 208 point mutations, 56 large indels (>50 bp), and 6 small indels (Fig.2; Supplementary Fig. 12-15; Supplementary Table 12). The mutations were consistently distributed across all eight lines and the five chromosomes (Extended Data Fig. 2).

**Fig. 2.**
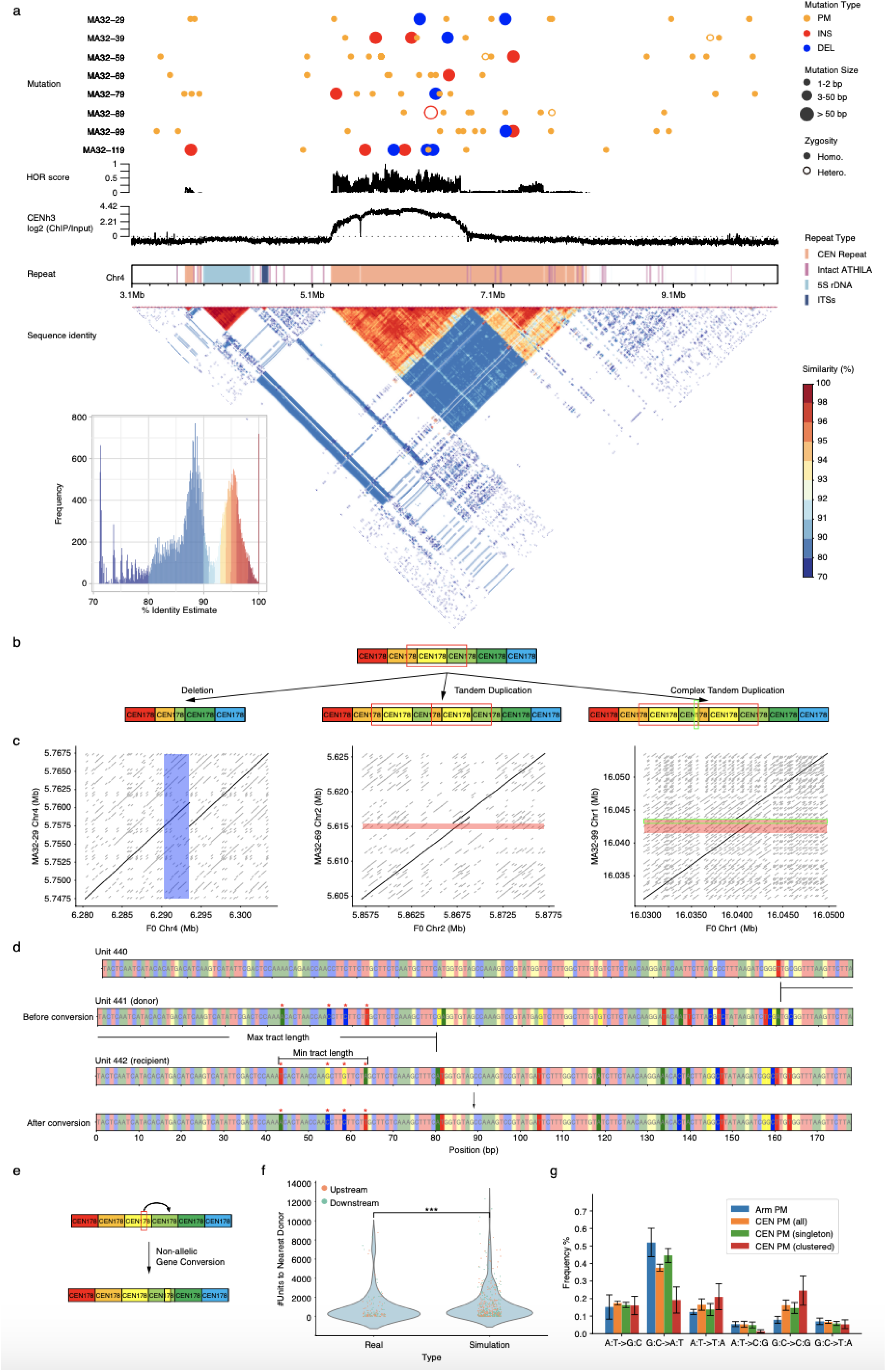
Mutations within centromeres. **a.** Eight rows of circles display mutation patterns in the centromere of chromosome 4 in MA32 samples, colour coded as point mutations (PMs; orange), insertions (red), and deletions (blue). Circle size reflects mutation length: small (1-2 bp), medium (3-50 bp), and large (>50 bp). Solid circles indicate homozygous mutations, while hollow circles indicate heterozygous ones. The two line plots below show higher-order repeat (HOR) scores and log2 CENH3 ChIP-seq enrichment^67^, respectively. Rectangles highlight repetitive regions, and the heatmap below represents pairwise sequence identity between non-overlapping 10 kb regions, with a histogram summarizing identity values in the lower-left corner. **b.** Schematics illustrate three mutation types: deletion (left), tandem duplication (middle), and complex tandem duplication (right), with red boxes highlighting mutated patterns. **c.** Dot plots show sequence identity between wild-type and mutated sequences. The longest identical sequences are shown in black and others in grey. Mutated regions are colour-coded: blue (deletion) and red (insertion). The green box highlights a mosaic region derived from nearby repeat units. **d.** A schematic example shows four point mutations introduced by a single non-allelic gene conversion (NAGC) event rather than four independent mutations. Colours represent nucleotide bases and asterisks indicate four point mutations located centrally within a single *CEN178* repeat unit. **e.** Schematic representation of NAGC in centromeres. The example shows the donor unit adjacent to the recipient, although in reality they may be separated. **f.** Violin plots compare distance to the donor unit (measured in repeat units) in observed point mutations and 500 random centromere positions. Donors are colour-coded as: downstream (blue) and upstream (pink), showing a significant difference (P = 2.9e-5, Mann-Whitney U test). **g.** Bar plots show the mutation spectra across different regions or mutation types: chromosome arms (blue), all centromeric point mutations (orange), singleton centromeric point mutations (green) and clustered centromeric point mutations (red). CEN, centromere; ITSs, interstitial telomeric sequences; MT, mitochondrial sequences; PM: point mutation; INS: insertion; DEL: deletion; TD: tandem duplication; CTD: complex tandem duplication.

The 56 large indels showed a striking pattern: all of them preserved the periodicity of the *CEN178* tandem repeats by adding or removing only complete repeat units (Fig. 2b, c), thereby maintaining the structural integrity of the satellite array. Insertions occurred more frequently (paired Wilcoxon signed-rank test, P = 0.062) and were significantly larger than deletions (Mann–Whitney U test, P = 0.0024): the 21 deletions removed 1 to 18 repeat units (avg. size: 923 bp), whereas the 35 insertions added 1 to 31 units (avg. size: 1870 bp) (Extended Data Fig. 3). In all cases, the inserted sequences were either identical to adjacent repeat units (tandem duplications) or composed of mosaics of several adjacent repeat units (complex tandem duplications) (Fig. 2b, c). In addition, many indels generated novel repeat units at their breakpoints by combining segments of two neighbouring units (Fig. 2b, c).

The 208 point mutations were found across the entire length of centromeres (incl. 94.7% that occurred in the *CEN178* repeat units and 5.3% that occurred in ATHILA elements). However, 59 (28.4%) of them clustered locally into 16 mutation groups (Methods). For example, four of the point mutations were clustered within a 20 bp region in the centre of a single *CEN178* repeat unit (Fig. 2d). The adjacent upstream repeat unit contained the exact same sequence variants as the ones introduced by the four mutations (Fig. 2d, e). For 15 of the 16 clusters, we found such putative donor sequences located on the same chromosome, at a median distance of 16 repeat units between the mutated cluster and the putative donor repeat unit.

These patterns are most parsimoniously explained by non-allelic gene conversion (NAGC) events that copies sequence variants between *CEN178* repeats. The average length of the converted regions was 58.5 bp (Supplementary Table 11), which is longer than the previously estimated tract lengths for non-crossover-associated gene conversions during meiosis (25-50 bp)^18^.

When searching for putative donor sequences of the 149 singleton point mutations (which could, in principle, also arise through NAGC events), we found such sequences for almost all, namely 129 (86.5%). We were concerned, however, that such patterns could also occur by chance: when simulating random point mutations, donor-like matches were observed in 86% of cases. While this suggested that the point mutations less frequently introduced by NAGCs, the distances between mutations and their putative donor sequences were significantly shorter in the real data than in the simulations (Fig. 2f), implying that at least a subset of the singleton point mutations also arises from NAGC events rather than by spontaneous mutations.

The existence of NAGC events was further supported by differences in the mutation spectrum. While GC→AT transitions are the most common type of point mutations in chromosome arms (∼50-60%)^14,15^, their frequency among point mutations in the centromere was significantly reduced to ∼40%. This difference could not be explained by a reduced GC content, as centromeres even have a slightly higher GC content (37.4%) as compared to chromosome arms (36.2%). Considering only clustered mutations (those that most likely result from NAGC events), the frequency of GC→AT transitions was even reduced to ∼20% (Fig. 2g). Interestingly, this transition rate reflects the existing sequence differences between repeat units and does not require any GC-biased processes during the conversion (Extended Data Fig. 4). Consistently, the frequency of GC→AT transitions among singleton point mutations (some of which might also be introduced by NAGC events) was ∼45%.

Taken together, centromeric regions show a strikingly distinct mutation spectrum compared to the unique regions of chromosome arms. Centromere mutations are characterized by periodicity-preserving indels spanning several kilobases that add or remove complete repeat units, which can lead to changes in the repeat arrays without disrupting repeat periodicity itself. In addition, non-allelic gene conversions, which are characterized by the conversion of short sequence tracts between adjacent repeats, contribute to a drastically increased mutation rate in the centromere.

### Mutation rate in the centromere

We estimated the point mutation rate in centromeric repeat arrays (including both spontaneous mutation and those arising from NAGCs) to be 6.5 × 10^-8^ (95% CI: 5.6–7.5 × 10^-8^) per site per generation. This is almost tenfold higher than recent estimates for chromosome arms in *A. thaliana*^15^ (Table 1). With the exception of methylated cytosines in transposable elements, which also exhibited elevated mutation rates, all other genomic regions show significantly lower point mutation rates^15^.

**Table 1.**
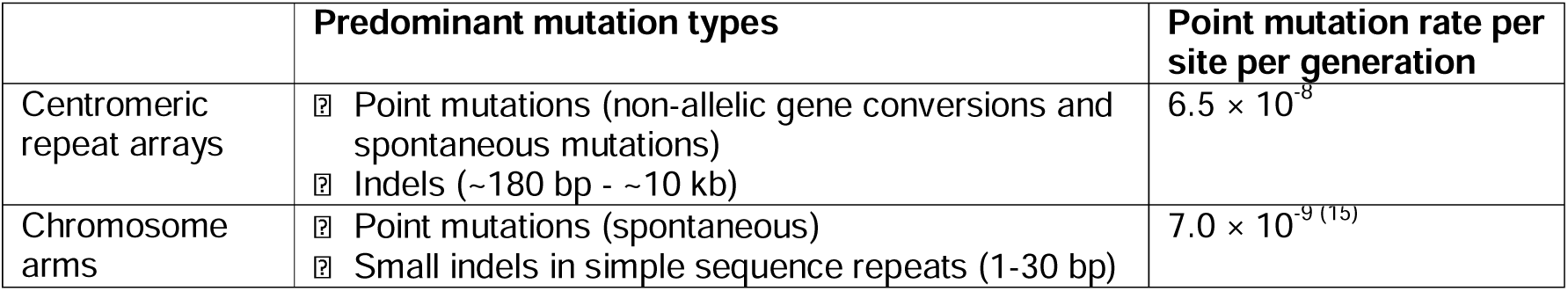
Different mutational dynamics in centromeres and chromosome arms.

To disentangle the contributions of the different mutational processes, we decomposed point mutations based on their observed differences in their transition/transversion (Ti/Tv) ratios (Methods). This analysis indicated that approximately 30.3% of point mutations in the centromeres arise from spontaneous mutations, whereas 69.7% of the point mutations are attributable to NAGC. This corresponds to a spontaneous mutation rate of 2.0 × 10^-8^ per site per generation, which is still approximately threefold higher than in chromosome arms and a NAGC-associated mutation rate of 4.5 × 10^-8^ per site per generation.

Taken together, these results indicate that centromeric point mutations are markedly elevated and are driven by both frequent non-allelic gene conversion events and spontaneous mutations.

### Centromeric TE movement is undetectable over recent timescales

Transposable elements (TEs) are common elements in centromeric repeat arrays. In Arabidopsis, ATHILA transposable elements are enriched in the centromeric and pericentromeric regions, and previous studies have suggested that these elements have a substantial contribution to the evolution and divergence of the centromeres^1,5^.

Across the eight MA32 lines, however, propagated for a total of 256 generations, we did not find a single TE insertion or deletion event in the centromeres. In addition, mutations in the TEs were underrepresented as compared to the rest of the centromere. Only 11 (5.3%) of the centromeric point mutations occurred within ATHILA elements, despite accounting for 12.4% of the centromeric sequence in the Col-0 genome.

To find footprints of recent TE activity in the centromere, we analysed the genomes of five Arabidopsis individuals of the HPG1 group, a homogeneous lineage of Arabidopsis that recently colonized North America. Most likely, a single ancestor of this group was introduced from Eurasia approximately 400 years ago^19–21^. The five HPG1 individuals were collected from geographically distinct locations and constitute a subset of the HPG1 group, which are non-admixed, non-recombined and quasi-identical individuals that can be differentiated only by the mutations that they accumulated since they separated^19^ (Fig. 3a; Supplementary Table 1). These genomes provide a “natural MA experiment” that allowed us to find mutations that accumulated for ∼1,500 generations (assuming an average generation time of 1.3 years)^19^.

**Fig. 3.**
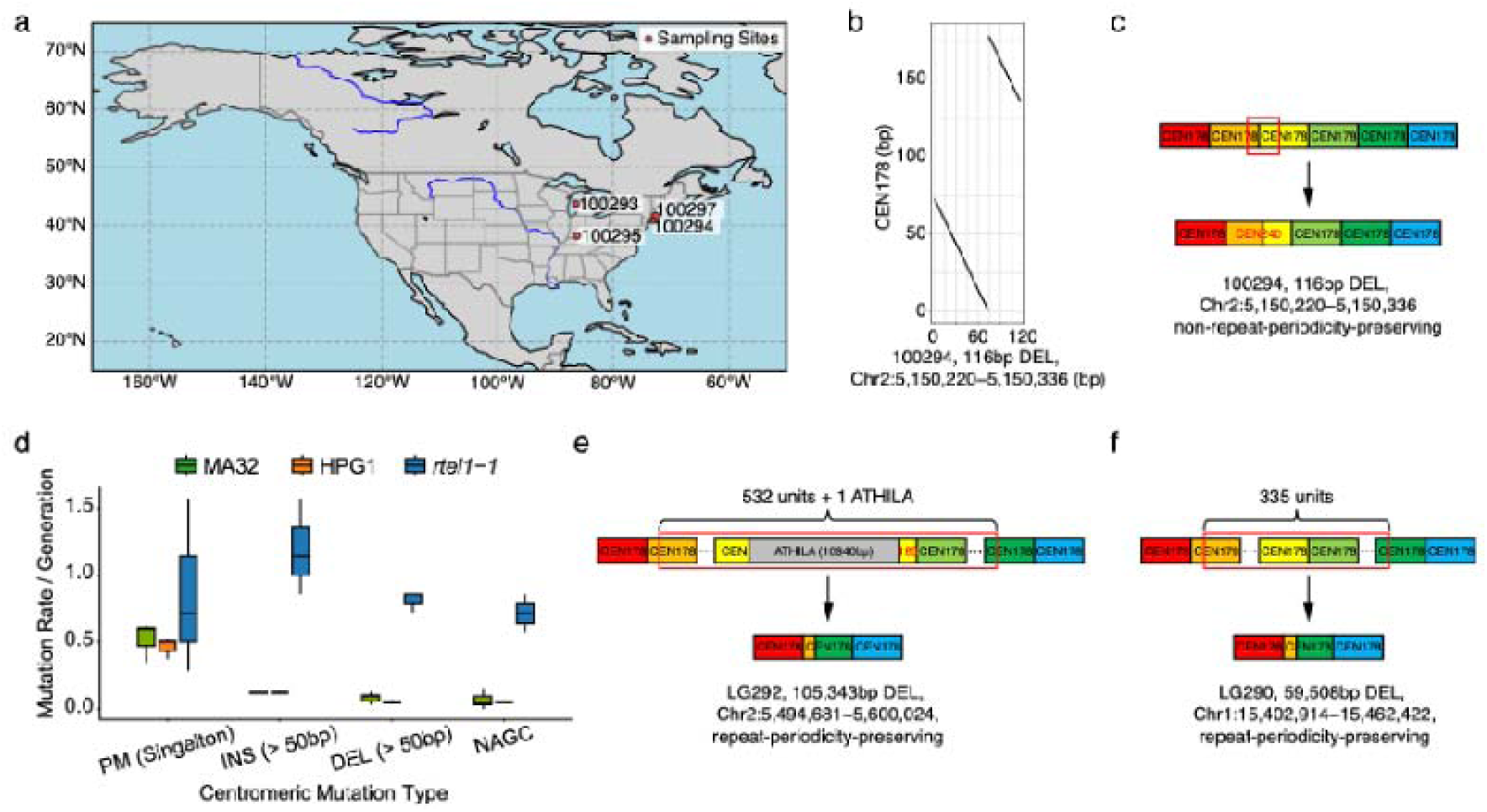
Accumulation of centromeric mutations in HPG1 lines and *rtel1-1* mutants. **a.** Geographic distribution of sampling locations for five HPG1 samples; no coordinate information is available for sample 100298. **b-c.** A 116 bp deletion in HPG1 that does not preserve repeat periodicity. **b,** Dot plot showing alignment of the 116 bp deletion to the *CEN178* consensus sequence. **c,** Schematic representation of the 116 bp deletion. **d.** Box plots showing the number of mutations per generation across three groups: MA32 (n = 8), HPG1 (n = 5), and *rtel1-1* (n = 3). Clustered point mutations are classified as NAGC events, whereas isolated point mutations are classified as singleton point mutations. Mann-Whitney U tests comparing the distributions of point mutations and indels between MA32 and HPG1 show no significant differences (*P* >0.05). **e-f.** Schematics of the two largest deletions identified in *rtel1-1*: 105,343 bp (**e**) and 59,508 bp (**f**). The former includes a complete ATHILA element. Despite their large sizes, both deletions preserve repeat periodicity. Breakpoint positions are shown to scale in the schematics.

We assembled the genomes of all five HPG1 accessions using PacBio long read sequencing. However, as in the MA lines, we found no evidence of TE movement in centromeres of the HPG1 genomes: all 152 intact ATHILA sites in these centromeres were conserved across all five individuals (Supplementary Fig. 16). In fact, we even did not find any evidence of TE insertions, deletions, or transposition events anywhere in the HPG1 genomes. This is in contrast to previous population-scale analyses of the TE diversity in 216 accessions that estimated that TE mutations arise at a rate of less than one insertion per 60 generations^22^, as well as with reports of individual accessions harbouring dozens of unique TE insertions^1^. The absence of TE insertions or deletions in the HPG1 accessions, which have accumulated mutations over ∼1,500 generations, suggests that TE insertion events may not occur at a constant rate but instead in occasional bursts^23^, such that no events are observed over these extended time intervals.

Despite the absence of TE movement, we identified substantial sequence variation within the centromeric arrays of HPG1 accessions. In total, we detected 1,415 homozygous mutations, including 1,113 point mutations (28 located within ATHILA elements), 275 large, kb-scale indels (>50 bp, max: 13,695 bp), and 27 small indels (<50 bp) (Supplementary Table 13). Consistent with the patterns observed in the MA lines, kb-scale insertions were more frequent than deletions (189 insertions vs. 86 deletions) and almost all of these indels preserved the repeat periodicity, with a single exception: a 116 bp deletion disrupted the tandem repeats and introduced a unique 240 bp-fragment into the repeat array (Fig. 3b, c; Extended Data Fig. 3).

The observed number of point mutations in HPG1 centromeres is in close agreement with the empirically estimated mutation rate (Fig. 3d). In total, 367 of the 1,113 point mutations formed 81 clusters within centromeric regions. Of those, 61 clusters could be linked to putative donor sequences, consistent with NAGC events. The remaining clusters lacked identifiable donors, possibly due to subsequent mutations that have obscured sequence similarity between the donor and mutation cluster.

### Homology-directed repair drives centromeric mutation

A common feature of tandem-repeat-preserving indels and NAGC events is their sequence similarity to other regions within the centromeric repeat array. This suggests that homology-directed DNA repair might contribute not only to the stabilization of the repeat array but also to its mutagenesis, consistent with the known propensities of homology-directed repair to occur in regions with local sequence similarities and to occasionally introduce mutations^24^.

To directly test whether dysregulated homology–directed repair alters centromeric mutational dynamics, we analysed MA lines deficient in RTEL1 (REGULATOR OF TELOMERE ELONGATION 1), a DNA helicase that dismantles recombination intermediates such as D-loops and thereby limits inappropriate or prolonged homologous recombination^25^. RTEL1 is required for genome stability by processing replication-associated DNA structures and suppressing excessive homologous recombination, particularly in repetitive regions. In Arabidopsis, loss of RTEL1 not only leads to hyperrecombination^26^ but also to a massive reduction of 45S rDNA tandem repeats^27^. The use of *rtel1-1* mutant MA lines enabled us to assess how homology-directed DNA repair impacts centromeric mutation rates and spectra.

We assembled the genomes of three MA lines using PacBio derived from a *rtel1-1* mutant, which were propagated for seven generations. Overall, we found 150 homozygous mutations in the centromere, including 89 point mutations, 57 large indels (33 insertions, 24 deletions) and four small indels (Supplementary Table 14). As in wild-type centromeres, point mutations in *rtel1-1* centromeres show the strong tendency to form local clusters: with 71 of 89 mutations in 14 distinct clusters. Despite accumulating mutations only for 21 generations, the three *rtel1-1* MA lines carried more centromeric indels than the eight MA32 wild-type lines, which together accumulated mutations for 256 generations: the MA32 lines accumulated 0.2 indels and 0.1 NAGC clusters per generations while the *rtel1-1* mutants accumulated 2.0 indels and 0.7 NAGC clusters per generations (Fig. 3d).

Again, all large indels preserved the tandem repeat periodicity and also the size distribution of the centromeric indels in *rtel1-1* lines was similar to that in wild-type MA lines (Extended Data Fig. 3). However, two deletions in *rtel1-1* MA lines were markedly longer: 59,508 bp and 105,343 bp, removing 335 and 532 repeat units, respectively. Despite their size, both deletions preserved the tandem repeat periodicity intact (Fig. 3e, f), consistent with the role of RTEL1 in dismantling recombination intermediates, which otherwise can lead to large mutations.

Together, the *rtel1-1* mutation introduces a massive change in the mutation rate, but not the spectrum of centromeric mutations, which is consistent with RTEL1 acting as a safeguard of homology-directed DNA repair and thereby supports the involvement of homology-directed DNA repair in the mutational processes in the centromere also in wild-type plants. This nails down that homology-directed DNA repair is the major cause of centromeric sequence evolution.

Together, the *rtel1-1* mutation induces a massive increase in the centromeric mutation rate, while largely preserving the characteristic tandem-repeat-preserving spectrum observed in wild-type lines. This is consistent with RTEL1 acting as a safeguard against excessive homology-directed DNA repair, and supports the view that homology-directed repair makes a substantial contribution to centromeric mutational processes, including in wild-type plants.

### The emergence of homogenized blocks from centromere-specific mutations

In both animals and plants, the centromeric repeat arrays often contain large homogenized blocks composed of highly similar repeat units, including many HORs and spanning hundreds of kilobases to megabases^1,11^, which are typically distinct from surrounding repeat units (Fig. 4a). Often, the functional centromere is concentrated in highly homogenized blocks. But despite this conserved function, the homogenized blocks show little sequence conservation across the global population of Arabidopsis accessions (examples shown in Fig. 4b). Although several mechanisms have been proposed to explain their origin and turnover (e.g., layered expansion, tandem duplication, or unequal crossover^1,11^), how such homogenized blocks form remains unclear.

**Fig. 4.**
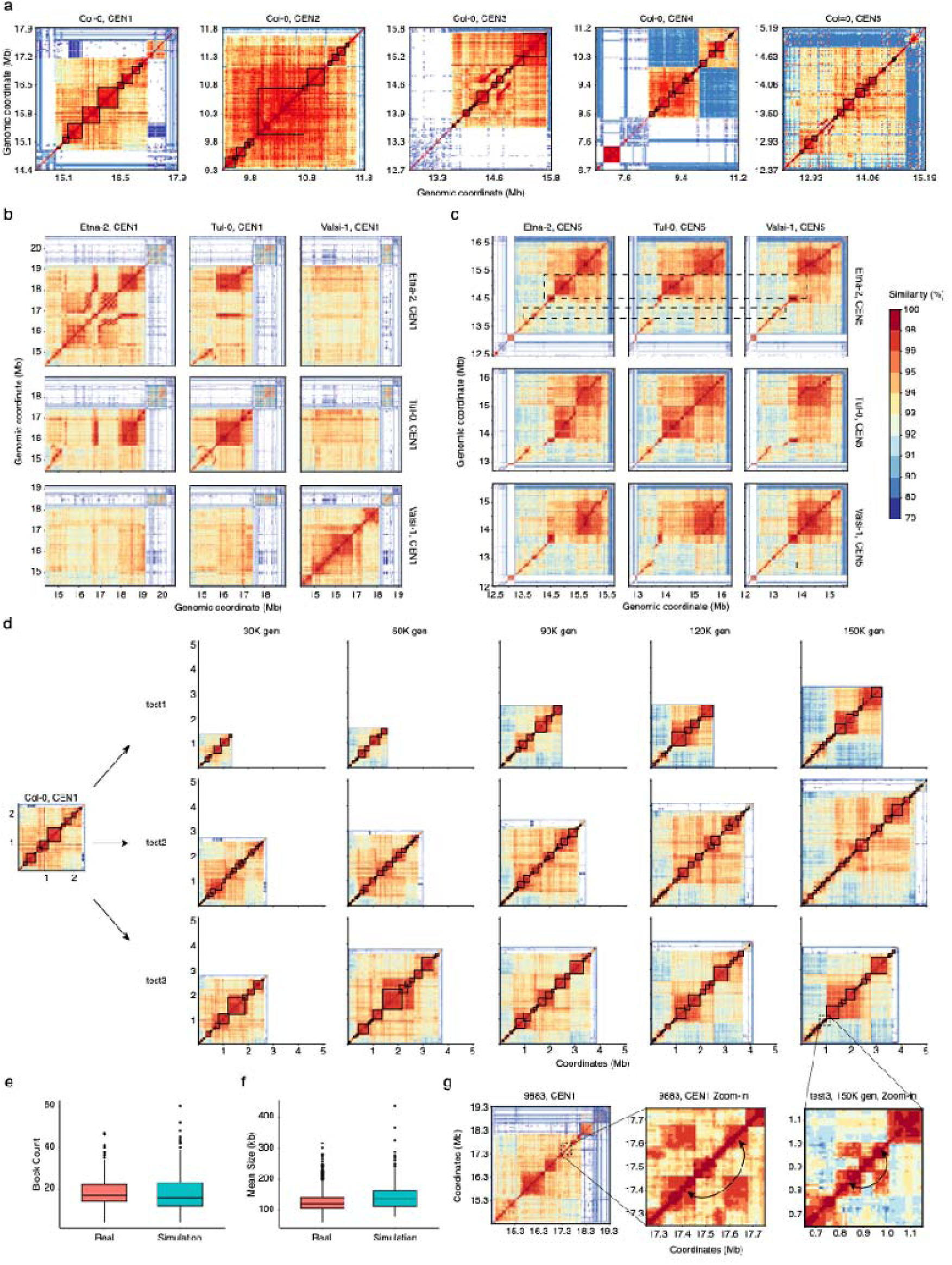
Simulations generate block structures resembling those of natural centromeres. **a.** Heatmap showing self-sequence identity across the centromeres of all five chromosomes in Col-0. Homogenized block structures are highlighted by black boxes (see Methods for definition). **b.** Heatmaps showing self- and pairwise sequence identity for the centromeres of chromosome 1 in Etna-2, Tul-0, and Valsi-1. Most centromeric sequences show limited conservation between accessions, similar to the relationships between Valsi-1 and Etna-2 or Tul-0. **c.** In contrast, accessions with similar centromeric haplotypes (as shown in centromeres of chromosome 5 in Etna-2, Tul-0, and Valsi-1) facilitate the detection of large-scale mutations, such as the putative deletion highlighted by the dashed box. **d.** Simulations based on observed mutation patterns in MA32, incorporating large deletions (average size 500 kb), were performed using Col-0 centromeric sequences for 15,000 generations. Changes in homogenized blocks (highlighted by black boxes) were observed over time. **e-f.** Box plots comparing the number (**e**) and mean length (**f**) of homogenized blocks between 570 real centromeres and 500 simulated centromeres (100 replicates per chromosome) at generation 150,000. **g.** Heatmaps showing examples of long-distance similarity in a real centromere (chromosome 1 of accession 9883) and in a simulated sequence (test3), with zoomed-in views of the highlighted regions. All heatmaps use a consistent colour scale representing sequence identity.

To investigate how the centromeric mutations contribute to the formation of homogenized blocks, we performed forward-in-time simulations in which the empirically estimated mutation spectrum was used to introduce realistic mutations into the centromeric sequences of all five Arabidopsis chromosomes over time. As the insertion rate was approximately twofold higher than the deletion rate, the simulated arrays expanded continuously, quickly reaching unrealistically large sizes (i.e., ∼20 Mb after ∼150,000 generations; Supplementary Fig. 17a).

This suggested that the observed mutation spectrum lacks balancing events that constrain the growth of the repeat array, similar to the very large deletions we observed in the *rtel1-1* mutants. Such events would likely be too rare to be observed in MA lines, and therefore should be large to have a substantial effect on array size. To identify such large-scale events, we searched for their footprints in 114 previously published genome assemblies of a global set of Arabidopsis accessions. By comparing all centromeric haplotypes against each other, we identified accessions sharing similar centromere haplotypes, which had not diverged too much, but enabled the detection of large-scale mutations (Method). For example, a ∼0.42-Mb region on chromosome 5 (13.80–14.22 Mb) in Etna-2 was absent in Tul-0, while the surrounding regions were structurally conserved, suggesting that this region was lost in Tul-0 (Fig. 4c). Likewise, another ∼0.64-Mb region (14.67–15.31 Mb) on the same chromosome in Etna-2 was absent in Valsi-1, where again the regions outside of the putatively deleted region were highly conserved (Fig. 4c; Extended Data Fig. 5 for additional examples).

We combined the empirically estimated mutation spectrum with such large-scale deletions (simulating sizes similar to the patterns observed in natural centromeres at a rate that stabilizes overall centromere size over time (3.0 × 10^-11^ per bp per generation; mean size: 0.5 Mb; Methods), corresponding to approximately one event every ∼2,800 generations). The simulations were initialized with the five centromeric sequences from the Arabidopsis reference genome, and mutations were sampled in each generation. Each simulation was run for 150,000 generations, consistent with the divergence time among major Arabidopsis population of approximately 120 - 90 kya^28^.

Within the simulated centromeres, existing homogenized blocks contracted and expanded, with some being lost and new blocks formed (Fig. 4d; see movies in Extended Data Files). Some of these new blocks eventually reached megabase sizes similar to those observed in natural centromeres. To be able to quantitatively compare such homogenized block patterns in the simulations to those of natural centromeres, we introduced a quantitative definition of the homogenized blocks that captures the visual patterns observed in centromeres while enabling objective comparison of well-defined homogenized blocks across different centromeres without relying on visual inspection alone (Fig. 4a; Methods).

After running the simulations, the resulting sequences recapitulated key features of the homogenised blocks of 570 natural centromeres, including similar block numbers and sizes (Fig. 4e, f). This illustrates how kilobase-scale indels can form megabase-scale homogenized blocks, while megabase-scale deletions are required to balance overall centromere sizes and thereby introduce severe changes to the global structures of the simulated centromeres.

The simulations also recapitulated long-distance similarities between distant blocks that can be observed in some natural centromeres (Fig. 4g; Extended Data Fig. 6)^5,29^. In the simulations, they arose when repeat units with distinct sequence variation started to expand to blocks and divided existing blocks into two distinct parts. The two separated parts then shared more similarity with each other than with the newly formed block in between. These patterns emerged exclusively through indel and point mutations and did not require long-distance recombination as previously proposed^5,29^.

Taken together, the simulations showed how centromeric mutations drive the turnover and formation of homogenised blocks, while rare large-scale deletions are required to balance the unequal frequencies of insertions and deletions to limit the overall size of the centromeres. The interplay of these processes is sufficient to generate centromeric repeat arrays that recapitulate key features of the blocks in natural centromeres, including block size and number.

## Discussion

In this study, we generated virtually error-free genome assemblies of *A. thaliana* mutation accumulation (MA) lines to investigate the mutational dynamics of centromeric tandem repeat arrays. Consistent with the high sequence divergence among natural centromeres, we observed an almost tenfold higher mutation rate in centromeres than in to chromosome arms, similar to rates reported for human centromeres and pericentromeric regions in Arabidopsis^11,15^.

Centromeric point mutations often occur in small clusters resembling sequence variation from nearby repeat units suggesting that a substantial fraction of these mutations arises from NAGCs, which reshuffle sequence variants among neighbouring repeats. In addition, frequent kilobase-scale indels almost exclusively preserved the tandem repeat periodicity, further supporting a homology-dependent mechanism underlying centromeric mutagenesis. Such processes could arise from the absence of meiotic crossovers^30,31^ or from homologous-driven DNA repair, where gene conversions and unequal crossing-overs have long been proposed to shape tandem repeat evolution through molecular drive and concerted evolution^2,32–38^. But also the observation of a non-repeat-unit-preserving deletion in the HPG1 lines, which accumulated mutations over ∼400 years, is consistent with the patterns found in natural centromeres, where most repeats are intact, but some rare truncated or partial repeats exist^1^.

To further investigate the mechanisms underlying centromere stability, we analysed the mutation dynamics in *rtel1-1* mutant MA lines. The *rtel1-1* mutants showed an unchanged mutational spectrum as compared to the spectrum in wildtype MA lines, yet the mutation rates of NAGC and tandem-repeat-preserving indels are markedly increased. This is supported by the role of RTEL1 to limit inappropriate or prolonged recombination and thus indicates that homology-directed repair plays a dual role in centromeric repeat arrays, both maintaining structural integrity and serving as a source of mutational change. The DNA helicase RTEL1 is involved in repair of replicative DNA damage in Arabidopsis^39^. In its wild-type form, RTEL1 dismantles DNA recombination intermediates, mainly by reversing aberrant annealing that might also occur during break-induced replication (BIR) or single strand annealing (SSA) reactions and would otherwise result in genome instabilities^40^.

Several features of the centromeric mutations are consistent with BIR^41–43^, which has been proposed to operate in centromeric regions, where one-ended double-strand breaks are repaired through homology-driven strand invasion and annealing reaction followed by conservative DNA synthesis^42,44^. These properties could explain the preservation of the repeat periodicity observed in centromeric indels. In addition, BIR is associated with a bias towards insertions and frequent template switching, providing a plausible mechanistic basis for the higher frequency of insertions relative to deletions and for the prevalence of complex tandem duplications in centromeric repeat arrays. Deletions might also arise by SSA between individual repeats^45^, which is enhanced in absence of RTEL1^46^. Indeed, also the analyses of natural centromeres revealed sequence patterns consistent with megabase-scale deletions, which are likely too rare to be captured by MA experiments.

Using forward-in-time simulations, we show that megabase-scale, homogenized blocks can arise through the accumulation of small, kb-scale mutations alone. However, as tandem-repeat-preserving insertions and deletions do not occur at the same frequency, additional, megabase-scale deletions are essential to balance overall size. While such events might be too rare to be identified in MA experiments, even very low frequencies of these large events are sufficient to balance centromere size.

Even though repeat homogenization can emerge independently of other centromeric features, this does not exclude the possibility that homogenization is influenced by or biased towards regions associated with CENH3 binding or kinetochore formation^1,47,48^. More broadly, these dynamics resemble concerted evolution, as described for rDNA and other satellite DNA families^31,48–51^, and are consistent with feedback models of satellite repeat evolution^52^, providing a mechanistic basis for the emergence of such patterns.

The simulations further suggest that changes in repeat unit consensus do not require the *de novo* formation of entire centromeric arrays but can arise through gradual, cumulative sequence changes^2,36,37^. Given that centromere function does not depend on specific repeat sequences, reduced meiotic recombination and relaxed selective constraints in these regions may facilitate the diversification and propagation of tandem repeat arrays, ultimately enabling the birth of new centromeric consensus sequences.

But there are still some unresolved observations. Though local, centromere-specific mutations shape the centromeric repeat arrays, the consensus sequences across the five chromosomes of Arabidopsis are similar (∼95% sequence identity between the chromosome-specific consensus sequences). This suggests inter-chromosomal exchange of individual repeats, potentially facilitated by centrophilic TEs, NAGCs or population dynamics^53^. However, we find no direct evidence for such exchange in our dataset. Despite analysing mutations accumulated over ∼1,500 generations, we did not detect any TE-associated mutations or inter-chromosomal NAGCs events.

Taken together, our study provides a comprehensive analysis of the spectrum and frequency of centromeric mutations in Arabidopsis, showing how point mutations, tandem-repeat-preserving indels together with rare large-scale deletions shape centromere structure over evolutionary timescales. Our findings offer new insights into long-standing questions about centromere evolution and provide a mechanistic framework for how these highly dynamic genomic regions are both maintained and remodelled.

## Online Methods

### Plant material

Four sets of *Arabidopsis thaliana* materials were analysed in this study, including MA16, MA32, HPG1 accessions, and the *rtel1-1* mutant (Supplementary Table 1). The MA16 lines (two lines, samples A and B) were derived from a trans-generational mutation accumulation (MA) experiment, originating from a single Col-0 mother plant (Nottingham Arabidopsis Stock Centre ID N1092) and propagated independently for 16 generations by self-pollination and single-seed descent (SSD). The MA32 dataset comprised eight lines were obtained from a previous MA experiment^54^ and propagated for 32 generations under a similar SSD scheme. The HPG1 dataset included five natural accessions representing natural populations that diverged approximately 400 years ago^19–21^ and were used to investigate recent centromeric transposable element activity. The *rtel1-1* mutant (three samples; SALK_113285) was propagated for seven generations by SSD to assess the role of homology-directed repair in centromere evolution.

### DNA extraction and PacBio sequencing

For MA16 samples A and B, HMW DNA was extracted from 1.5 g of pooled vegetative tissue using the NucleoBond HMW DNA kit (Macherey-Nagel). DNA quality was assessed using a FEMTOpulse system (Agilent), and concentration was measured with a Quantus fluorometer (Promega). HiFi SMRTbell libraries were prepared using the SMRTbell® Express Template Prep Kit 2.0 (PacBio), including fragmentation with g-TUBEs (Covaris) and size selection using SageELF (Sage Science). Libraries were sequenced on the PacBio Sequel II platform at the Max Planck Genome Centre (MP-GC), Cologne, Germany. In addition, PCR-free Illumina paired-end libraries were prepared from independently extracted DNA (Macherey-Nagel DNA Maxi kit) and sequenced by Novogene.

For eight MA32 lines, HMW DNA was extracted at the Max Planck Institute for Biology Tübingen using a modified protocol^9^, including β-mercaptoethanol during lysis and a phenol purification step. DNA was further purified using two rounds of bead cleanup (SeraMag SpeedBeads and AMPure PB beads). Libraries were prepared using the SMRTbell® prep kit 3.0. HiFi sequencing was performed on the PacBio Revio device at MP-GC.

For five HPG1 accessions, HMW DNA extraction and library preparation were performed at the Max Planck Institute for Biology Tübingen using the same protocol as for MA32. Libraries were prepared with the HiFi SMRTbell® Express Template Prep Kit 2.0 (PacBio). Libraries were size-selected using the BluePippin system (Sage Science) and sequenced on a Sequel II system with Binding Kit 2.2 at the Max Planck Institute for Biology Tübingen.

For three *rtel1-1* samples, HMW DNA was extracted using a kit-based protocol at KIT (Karlsruhe Institute of Technology), followed by HiFi library preparation using the SMRTbell® prep kit 3.0 and libraries sized with BluePippin (Sage Science). Sequencing was performed on the PacBio Revio device at MP-GC. In addition, PCR-free Illumina paired-end sequencing was carried out at the Institute of Clinical Molecular Biology (IKMB), Kiel.

### Nanopore sequencing and basecalling

For the two MA16 lines and the eight additional MA32 lines, library preparation was performed using the Ligation Sequencing gDNA – Native Barcoding Kit 24 V14 (SQK-NBD114.24, Oxford Nanopore Technologies). The resulting libraries were loaded onto FLO-PRO114M flow cells, and sequencing was conducted on a PromethION 2 Solo platform.

ONT sequencing data were basecalled with Dorado v1.1.1 (https://github.com/nanoporetech/dorado/) using the sup model for high accuracy basecalling, including modified base detection (5mC and 5hmC) and move table output. Barcode demultiplexing was guided by the SQK-NBD114-24 kit and a sample sheet. Resulting BAM files were split by barcode using samtools^55^ split v1.17 for downstream analysis.

### Genome assembly and scaffolding

We tested multiple assemblers^56–58^ to assemble the A1 genome and found that Hifiasm^56^ v0.16.0-r3699 produced the most contiguous assembly. We used Hifiasm^56^ with the parameter "-l0" for all four generation-16 samples (A1, A2, B1 and B2), eight generation-32 MA lines and *rtel1-1* mutant. Organellar contigs were then identified based on sequence alignment to TAIR10^59^ mitochondria and chloroplast references (GCF_000001735.4), retaining those with ≥80% identity and coverage. Non-organellar contigs were scaffolded into pseudo-chromosomes using RagTag^60^ v1.0.1, based on alignment to the Col-CEN^5^ reference genome. We further evaluated polishing strategies for HiFi-based assemblies using the MA16 lines. Polishing introduced over-corrections and led to an increase in assembly errors (Supplementary Results). Therefore, no polishing was applied to HiFi-based assemblies in subsequent analyses. Additionally, genome assemblies for five HPG1 samples were generated using Hifiasm^56^ v0.16.1-r375 and scaffolded with RagTag^60^ v2.0.1 (scaffold -q 60 -f 30000 -I 0.5 -remove-small), excluding contigs <100 kb.

To complement the HiFi-based assemblies, ONT long-read assemblies were also generated using Hifiasm^61^ v0.25.0 with the parameters --ont -l0 --rl-cut 10000 --sc-cut 15, which restricts assembly to high-quality reads ≥10 kb with estimated QV ≥15. The ONT contigs were filtered to remove organellar sequences and further polished. ONT reads were firstly aligned to the contig-level assemblies using the Dorado aligner, followed by sorting and indexing with samtools^55^ v1.19.2. Coverage profiles were computed using mosdepth^62^ v0.3.1, and regions with abnormal coverage were filtered prior to polishing using a custom script. To evaluate polishing strategies, two approaches were tested for the MA16 lines: polishing with move table information (“with moves”) and without move table information. Comparative assessment showed that polishing with move table information resulted in fewer assembly errors. Therefore, all subsequent polishing of ONT-based assemblies was performed using the move-aware Dorado polishing mode with region filtering and GPU acceleration.

The polished contigs were then scaffolded with RagTag^60^ using the Col-CEN^5^ reference, following the same strategy as for the HiFi assemblies.

### Assembly evaluation

We computed Benchmarking Universal Single-Copy Ortholog (BUSCO) scores using BUSCO^63^ v5.2.2 with the parameters “-l embryophyte_odb10 -m genome.” Additionally, we assessed consensus quality and completeness using Merqury^13^ v1.3 by comparing *k*-mers in the *de novo* assemblies with those from Illumina short reads. *K*-mer databases (k=18) were generated for each Illumina paired-end read using Meryl^13^ v1.3 and then merged with Meryl’s union-sum function. Merqury^13^ was subsequently applied to each assembly to obtain genome-wide consensus quality values (QV) and completeness scores.

### Repeat annotation

Following the approach of Rabanal et al. (2022)^9^, we used RepeatMasker v4.0.9 (http://www.repeatmasker.org) with a custom library (-lib rDNA_NaishCEN_telomeres.fa -nolow -gff -xsmall -cutoff 200) to annotate 5S rDNA, 45S rDNA, and telomere sequences. Mitochondrial insertions on Chr2 were identified by aligning to the TAIR10^59^ mitochondrial sequence using minimap2^64^ v2.24-r1122. Centromeres were annotated using TRASH^65^ v1.2 (--seqt CEN178.csv --horclass CEN178 --par 5), and the HOR score of each centromeric repeat unit was calculated. For samples A and B, we further annotated simple sequence repeats with a custom Python script to identify mono-, di-, tri-nucleotide and hepta-repeats. Bedtools^66^ v2.29.0 intersect was used to compare assembly errors and mutations across different repeat types.

We further identified the CENH3 enrichment regions. Raw paired-end CENH3 ChIP–seq (SRR4430537) and corresponding input reads (SRR4430555)^67^ were firstly adapter-trimmed and quality-filtered using Cutadapt^68^ v5.1, and aligned to the reference genome using Bowtie2^69^ v2.5.1 with sensitive parameters “--very-sensitive --no-mixed --no-discordant”. Alignments were sorted and indexed using Samtools^55^ v1.19.2, and genome-wide coverage tracks were generated using deepTools^70^ v3.5.6 with BPM normalization. Enrichment of ChIP signal over input was calculated as log2 ratios using bigwigCompare.

### Identification of assembly errors

To identify assembly discrepancies between replicate genome assemblies, we performed pairwise alignments of the six assemblies (A1-A3, B1-B3) using minimap2^64^ v2.24-r1122 (-ax asm5 --eqx). Assembly differences were identified with SYRI^71^ v1.0. To determine which replicate contained the assembly error, we mapped both HiFi and Illumina reads to the reference and their corresponding assemblies using minimap2^64^ v2.24-r1122 (-ax map-hifi) and BWA-MEM^72^ v0.7.17-r1188, respectively. Alignments were sorted and converted to BAM files with Samtools^55^ v1.19.2. Additionally, HiFi reads from each replicate were aligned to the opposing replicate’s assembly for cross-comparison. For regions flagged by SYRI^71^, we examined alignments in IGV^73^ v2.13.0. If HiFi and Illumina data aligned cleanly to one assembly but showed mismatches in the other, the error was attributed to the mismatching assembly, and the error-free one was considered correct. This approach allowed us to systematically identify and verify which sample contained assembly errors. It is worth noting that regions near GA repeats exhibited incomplete assemblies due to reduced HiFi read coverage^9^. These were classified as incomplete rather than erroneous assemblies, as the low-depth pattern was consistent across samples.

### Identification and validation of mutations

To identify mutations for each of the separate groups, MA16, MA32, HPG1, *rtel1-1*, we first selected the most contiguous assembly and then aligned each of the other assembly with minimap2^64^ v2.24-r1122. Alignments were sorted with Samtools^55^, and sequence differences were detected using SYRI^71^. To validate candidate mutations and determine their zygosity, HiFi, ONT, and Illumina reads were aligned to both the corresponding sample assembly and the reference assembly (long reads using minimap2^64^ v2.24-r1122 and short reads using BWA-MEM^72^ v0.7.17-r1188). Mutations were visually assessed in IGV^73^ to confirm their presence and to assess whether they are homozygous or heterozygous based on read alignment. To distinguish mutated from wild-type alleles, we referenced the TAIR10^59^ and the Col-CEN^5^ assemblies, assigning the allele matching the reference as wild-type and the differing one as a mutation. Variants shared by at least two individuals within a group were considered likely derived from segregation of pre-existing heterozygous variants in the parental material and were excluded from downstream analysis.

For MA16 samples, mutations were validated genome-wide, whereas for the MA32 lines, HPG1 samples, and *rtel1-1* mutants, analyses focused primarily on centromeric regions (defined as the interval spanning the outermost *CEN178* repeats and ATHILA elements), as well as single-nucleotide variants located on chromosome arms. Clusters of mutations were defined as groups of variants with consistent zygosity located within 1 kb of each other; variants separated by ≤1 kb were considered part of the same mutation cluster.

To ensure accurate coordinate mapping and facilitate identification of potential non-allelic gene conversion (NAGC) donor sequences, ancestral (F0) reference assemblies were reconstructed. Specifically, positions identified as mutations relative to the reference but shared across other samples were reverted to the wild-type allele using consensus function implemented in bcftools^55^ v1.16. All alignments and mutation validations were subsequently repeated against these reconstructed reference assemblies.

### Curation of mutation calls

Due to the highly repetitive nature of centromeric sequences, alignment errors can introduce fragment a single large variant into multiple smaller calls. To further correct for alignment artefacts and refine mutation calls, we developed a word-based alignment approach. For each candidate mutation, the mutated sequence and the corresponding reference sequence, each including 10 kb flanking regions, were extracted using seqtk v1.4-r122 subseq function (https://github.com/lh3/seqtk). Exact matches were identified using a custom Python script based on k-mer indexing (k = 150, no mismatches), using long exact matches as anchors between sequences in both forward and reverse-complement orientations. This approach identifies long, uninterrupted matching segments as high-confidence anchors, avoiding mismatches and gaps that can confound conventional alignment algorithms in repetitive regions. Based on these anchors, mutations were redefined using a left-alignment strategy to obtain the most parsimonious representation of each variant. For precise characterization of large indels, the word-based alignments were further visualized using dot plots generated by a custom Python script. Compared to standard alignment methods, this mismatch-free approach enables more accurate delineation of mutation boundaries and reduces false-positive fragmentation of variants in centromeric regions.

### Assessment of repeat periodicity in centromeric indels

To assess whether large centromeric indel mutations preserve the periodic structure of the centromeric repeat unit, indel sequences were aligned to the *CEN178* consensus sequence using MUMmer^74^ v4.0.0beta2 (nucmer, --maxmatch -l 10 -c 20). Alignments were converted from .delta to .coords format using show-coords (-c), and dot plots were generated using the R package dotPlotly (https://github.com/tpoorten/dotPlotly) (-m 10 -q 10 -k 5 -l -x).

### Statistical assessment of mutation distribution

The curated mutation counts for each mutation type across samples and across chromosomes were analysed using chi-square goodness-of-fit tests to assess deviations from uniformity. For chromosome-level analyses, expected counts were scaled by centromeric sequence size to account for differences in mutation opportunity. P-values were adjusted using the Benjamini–Hochberg false discovery rate (FDR) method. Due to low counts and violation of test assumptions, insertions and deletions were combined into a single “indel” category to improve statistical power. Mutation distributions along chromosomes were visualized using the R package karyoploteR^75^.

### Identification of candidate donor sequences for NAGC

To identify potential donor sequences underlying non-allelic gene conversion (NAGC) events, we performed targeted searches within centromeric repeat arrays using mutation-containing sequences as queries.

For mutation clusters, the full mutated sequence spanning each cluster was used as the query to identify exact matches within centromeric repeats. Candidate donor sites were required to map to the same relative position within the repeat consensus as the mutation cluster. For singleton point mutations, all possible 20-bp sequences spanning each mutation site were extracted and used as queries, applying the same positional constraint within the repeat consensus.

When multiple candidate donors were identified, the repeat unit with the shortest distance to the mutation site (measured in repeat units) was selected as the most likely donor. In cases where only a single candidate donor was identified, it was always located on the same chromosome as the mutation. No instances were observed in which candidate donors were exclusively located on different chromosomes, suggesting that inter-chromosomal NAGC events are rare.

### Identification of mutations associated with novel TE insertions

Transposable elements (TEs) and ATHILA elements were annotated in both the reference and query genomes using EDTA^76^ and ATHILAfinder^77^, respectively. We used a TE presence/absence–based strategy to identify structural variants potentially associated with novel TE insertions.

Mutations of at least 50 bp between the reference and query genomes were previously identified using SyRI^71^. The coordinates of these SVs in both genomes were intersected with the corresponding intact TE annotations using bedtools^66^ intersect. An SV was considered to be associated with a novel TE insertion only if it overlapped a TE annotation in the query genome but showed no overlap with any TE annotation at the corresponding location in the reference genome.

To further validate the integrity and conservation of the identified ATHILA elements and to exclude the possibility of misidentification, we performed a phylogenetic reconstruction. For each identified ATHILA locus, sequences from five HPG1 samples were extracted and aligned using MAFFT^78^ (v7.407) with the --autoparameter. The resulting multiple sequence alignments were used to infer maximum-likelihood phylogenetic trees using FastTree (v2.1.11, https://github.com/rambaut/figtree) with the -nt (nucleotide) model. The resulting phylogenetic trees were visualized, annotated, and refined using iTOL^79^ (Interactive Tree Of Life, v7). The robust clustering pattern demonstrates that these ATHILA elements are orthologous across the HPG1 samples and have been maintained with high sequence integrity. These results provide compelling evidence that these elements were inherited from the ancestral genome and that no novel ATHILA insertion events or associated large-scale structural mutations have occurred at these loci across the sampled accessions.

### Mutation rate calculations

Mutation rates were calculated as mutations per nucleotide per generation using the formula:

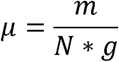

where *m* = total observed mutations, *N* = genome size (nucleotides), and *g* = generations. Mutations from all eight MA32 lines were pooled to calculate the overall mutation rate. Total mutations were divided by the cumulative nucleotide-generations (∑[*N* ⋅ *g*] = 3.032 × 100), yielding:

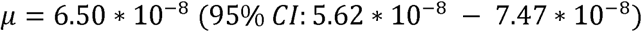

Confidence intervals (95%) were derived from Poisson statistics using the chi-square approximation:

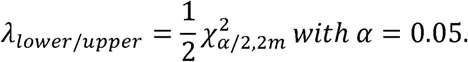

To distinguish spontaneous point mutation from those arising through NAGC, we decomposed the observed transition/transversion (Ti/Tv) ratios. Clustered mutations, which are characteristic of NAGC, exhibited a Ti/Tv ratio of 0.78, whereas mutations on chromosome arms (representing spontaneous mutations) showed a Ti/Tv ratio of 2.24. The overall Ti/Tv ratio of centromeric point mutations was 1.22. Based on previous studies suggesting that the Ti/Tv ratio of spontaneous mutations is similar between centromeres and chromosome arms^15^, we decomposed the observed overall Ti/Tv (1.22) into a linear combination of spontaneous mutations and NAGC-derived mutations. We estimated that 30.3% of point mutations arose from spontaneous mutations and 69.7% from NAGC. This yielded a spontaneous mutation rate of 2.0 × 10^-8^ per bp per generation and a NAGC-associated mutation rate of 4.5 × 10^-8^ per site per generation.

### Analysis of natural centromeres

To analyse the sequence features and structural patterns of natural centromeres, we used 114 publicly available high-quality HiFi-based genome assemblies of *Arabidopsis thaliana*, including 66 from Wlodzimierz *et al.* (2023)^1^ and 48 from Lian *et al.* (2024)^80^. Centromeric sequences were identified using both CentroAnno^81^ v1.0.2 and TRASH^65^ v1.2. Downstream analyses were primarily based on CentroAnno annotations due to its higher computational efficiency and scalability for large datasets. TRASH^65^ was additionally applied to ensure consistency with analyses performed on MA lines and to validate the robustness of centromere annotations across methods, which showed overall concordant results. Using the subseq function in seqtk v1.4-r122 (https://github.com/lh3/seqtk), we extracted the genomic regions spanning between the two outermost centromeric repeats for each chromosome. A total of 570 centromeric sequences (114 accessions × 5 chromosomes) were analysed. Sequence identity was quantified using ModDotPlot^82^ v0.9.8 for both self-comparisons and pairwise comparisons (restricted to the same chromosome across accessions), with parameters “-w 10000 -d 0”. It partitioned genomic sequences into non-overlapping windows of 10 kb and calculated similarity between all window pairs. Finally, sequence identity matrices were visualized as heatmaps using custom Python scripts.

### Formal definition of homogenized blocks

To formally describe the appearance of homogenized blocks, we developed a definition based on the pairwise similarity matrix from ModDotPlot^82^ v0.9.8. Homogenized blocks were defined as continuous genomic regions spanning at least five windows (≥50 kb), with a minimum average pairwise similarity of 97%. Self-comparisons and immediately adjacent windows were excluded from the calculation to avoid inflated similarity. For each region, the average similarity was computed across all valid window pairs. Nested blocks (i.e., overlapping or fully contained regions representing redundant detection) were subsequently filtered to retain only the largest non-redundant regions. Blocks were further filtered based on their overlap with centromeric repeat annotations, requiring at least 90% of each block to be annotated as centromeric repeat sequence, while blocks not meeting this criterion were excluded. For visualization, similarity matrices were plotted as heatmaps using custom colour palettes, and detected blocks were overlaid as rectangular annotations.

### Forward-in-time simulation of centromere evolution

We estimated mutation rates based on 270 homozygous mutations identified across eight MA32 samples, including 208 point mutations, 56 large indels, and 6 small indels. This corresponded to mutation rates of 6.5 × 10^-8^ per bp per generation for point mutations, 1.2 × 10^-8^ for large insertions (mean size 1,870 bp; 10.51 repeat units), and 6.9 × 10^-9^ for large deletions (mean size 923.1 bp; 5.19 repeat units).

To assess whether the observed mutation spectrum of clustered centromeric point mutations can be explained by NAGC and to exclude the contribution of GC-biased processes, we performed NAGC-only simulations under the same parameter framework. Five centromeric sequences were independently simulated, each with three replicates. In each replicate, NAGC events were iteratively introduced, and mutation spectra were calculated from the resulting sequences and compared to the observed clustered mutation spectrum. In addition, simulations were performed with varying recipient unit distances (adjacent, and separated by 1, 9, or 99 intervening repeat units) to evaluate the effect of spatial separation on mutation patterns. This analysis allowed us to determine whether NAGC alone can recapitulate the observed mutation spectrum without invoking additional mutational biases such as GC-biased gene conversion.

To estimate the frequency of NAGC events, we simulated gene conversion on the reference genome. Parameter exploration indicated that tract length primarily determines the number of introduced variants, whereas donor distance has minimal effect (Extended Data Fig. 4; Supplementary Fig. 18). Based on observed mutation clusters, NAGC tracts were modelled using a geometric distribution (mean = 0.05), with donors restricted to adjacent repeat units. Across 1,000 simulations per chromosome (five centromeres total), 6,684 point mutations were introduced, corresponding to an average of 1.34 point mutations per event and an estimated NAGC rate of 3.4 × 10^-^0 per bp per generation.

Using these estimated rates, we simulated centromeric repeat evolution on the five Col-0 centromeric repeats, incorporating spontaneous point mutations, NAGC events, and large insertions and deletions (schematic simulation pipeline in Supplementary Fig. 19). Each centromere was simulated independently for 150,000 generations with 20 replicates. These simulations revealed exponential array expansion due to a bias toward more frequent and larger insertions (Supplementary Fig. 17a), resulting in unrealistically large arrays (>20 Mb).

To balance array size, we introduced rare but large deletions, motivated by patterns observed across natural centromeres. Large deletions were modelled with an average size of 500 kb (2,809 repeat units) and a rate of 3.04 × 100¹¹ per bp per generation (to balance the array size). With this parameterization, each centromere was simulated for 100 replicates over 150,000 generations.

To visualize array dynamics, simulation outputs were plotted every 1,000 generations and compiled into movies using FFmpeg^83^ v7.0.1 (10 frames per second, libx264 encoding). To compare simulated and natural centromeres, self- and pairwise similarity analyses were performed using ModDotPlot^82^ v0.9.8, and homogenized blocks were identified from self-similarity matrices as described above.

## Supporting information

Supplementary Results

## Data availability

The HiFi reads for A1 used in this study were previously published in the context of the first assembly of the Arabidopsis centromeres^5^. Genome assemblies and PacBio sequencing data of three HPG1 samples (100293-100295) were previously analysed^84^. CENH3 ChIP–seq data (SRR4430537) and corresponding input data (SRR4430555)^67^ were retrieved from the NCBI Sequence Read Archive. To ensure comprehensive data access and support further analysis, we are releasing the complete set of HiFi and Illumina reads used in this study. Data related to the replicated MA16 genome assemblies are available through NCBI under BioProject accession numbers PRJNA1259971; the data for the eight MA32 assemblies can be found at PRJEB112076; the data for the HPG1 and *rtel1-1* assemblies can be found at PRJEB75768 and PRJNA1458610, respectively.

## Code availability

Scripts used in this article are available at https://github.com/schneebergerlab/replicated-assemblies-centromere-study.

## Acknowledgements

The authors would like to thank Craig Dent and Leon Rauschning (MPI-PZ) for helpful discussions and Katrin Fritschi (MPI-Biol) for technical help. This work was funded by the Deutsche Forschungsgemeinschaft (DFG, German Research Foundation) under Germany’s Excellence Strategy – EXC 2048/1 – 390686111 (KS) and the Transregio Collaborative Research Center TRR356/1 2023 ‘Genetic diversity shaping biotic interactions of plants’ (491090170) (DW, KS), the Max Planck Society (DW), the European Research Council (ERC) grant “BYTE2BITE” (101124694) (KS), the BBSRC (grant BB/W015250/1) (JT, LMS) and “Prime-A-Plant” (309944) (JT).

## Author contributions

XD, HP, DW and KS developed and supervised the project. LG, FR, SA, JAC, YT, JT, LMS, BH performed plant work and/or generated data. XD, FR, YT assembled the genomes. XD performed the data analysis with help from WBJ, FR and MTP. XD and KS wrote the manuscript with input from all authors. All authors read and approved the final manuscript.

## Figures

**Extended Data Fig. 1.**
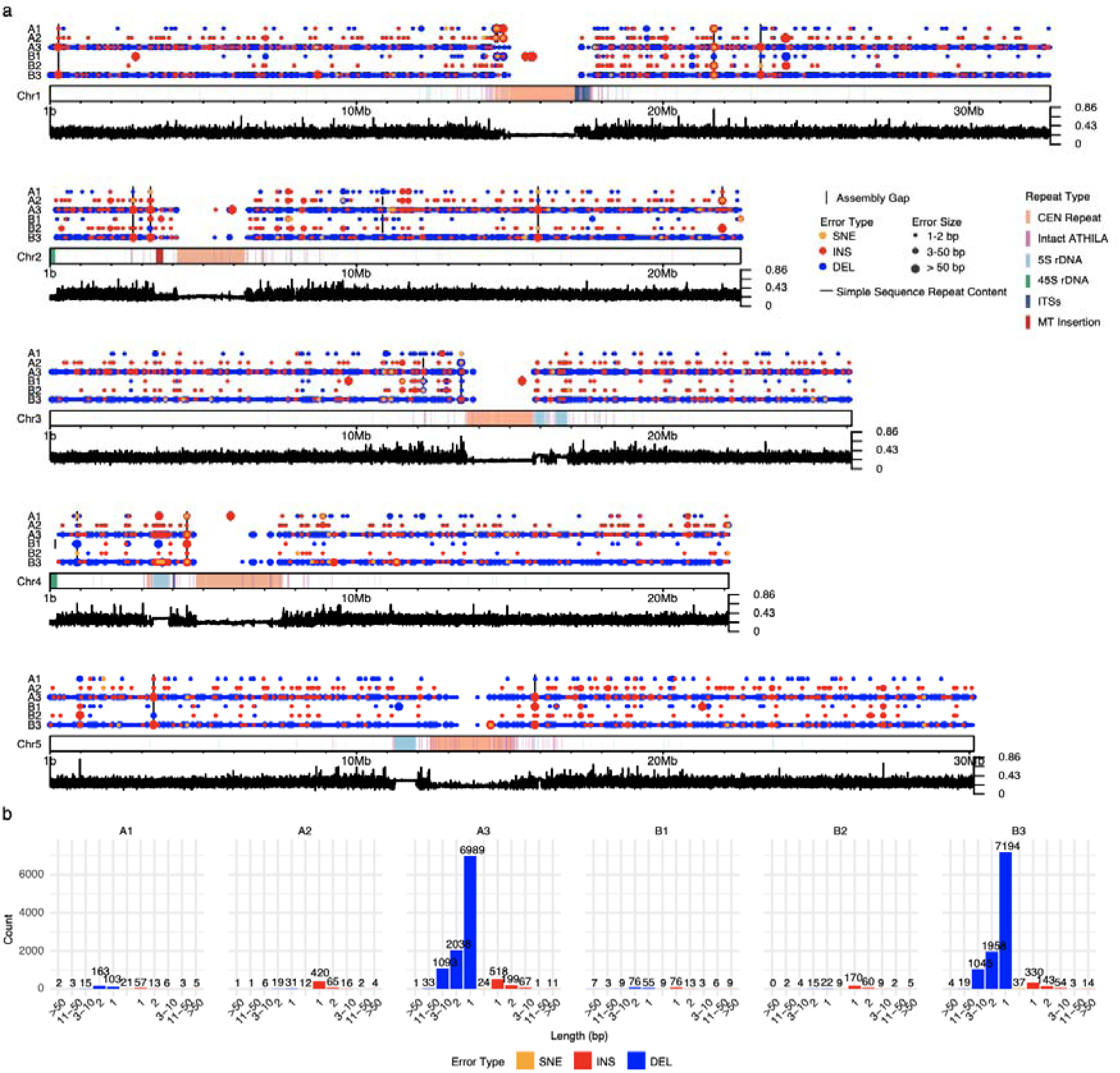
Assembly errors identified from long-read-based genome assemblies. **a.** Distribution of assembly errors across the five chromosomes. From top to bottom, tracks correspond to three independent sequencing and assembly replicates for samples A and B. A1–A2 and B1–B2 were generated using PacBio HiFi reads, whereas A3 and B3 were generated using Oxford Nanopore (ONT) reads. Each circle represents an assembly error, colour-coded by type: single-nucleotide errors (yellow), insertions (red), and deletions (blue). Circle size reflects error size: small (1-2 bp), medium (3-50 bp), and large (>50 bp). Vertical black lines indicate assembly gaps. Rectangles highlight repetitive regions: centromeres (peach), intact ATHILA elements (magenta), 5S rDNA (light blue), 45S rDNA (green), interstitial telomeric sequences (dark blue), and mitochondrial insertions (red). The bottom track shows the content of simple sequence repeats within 10 kb windows. **b.** Bar plots showing the distribution of assembly errors of different sizes across the six genome assem blies. CEN, centromere; ITSs, interstitial telomeric sequences; MT, mitochondrial sequences; SNE, single-nucleotide error; INS, insertion; DEL, deletion.

**Extended Data Fig. 2.**
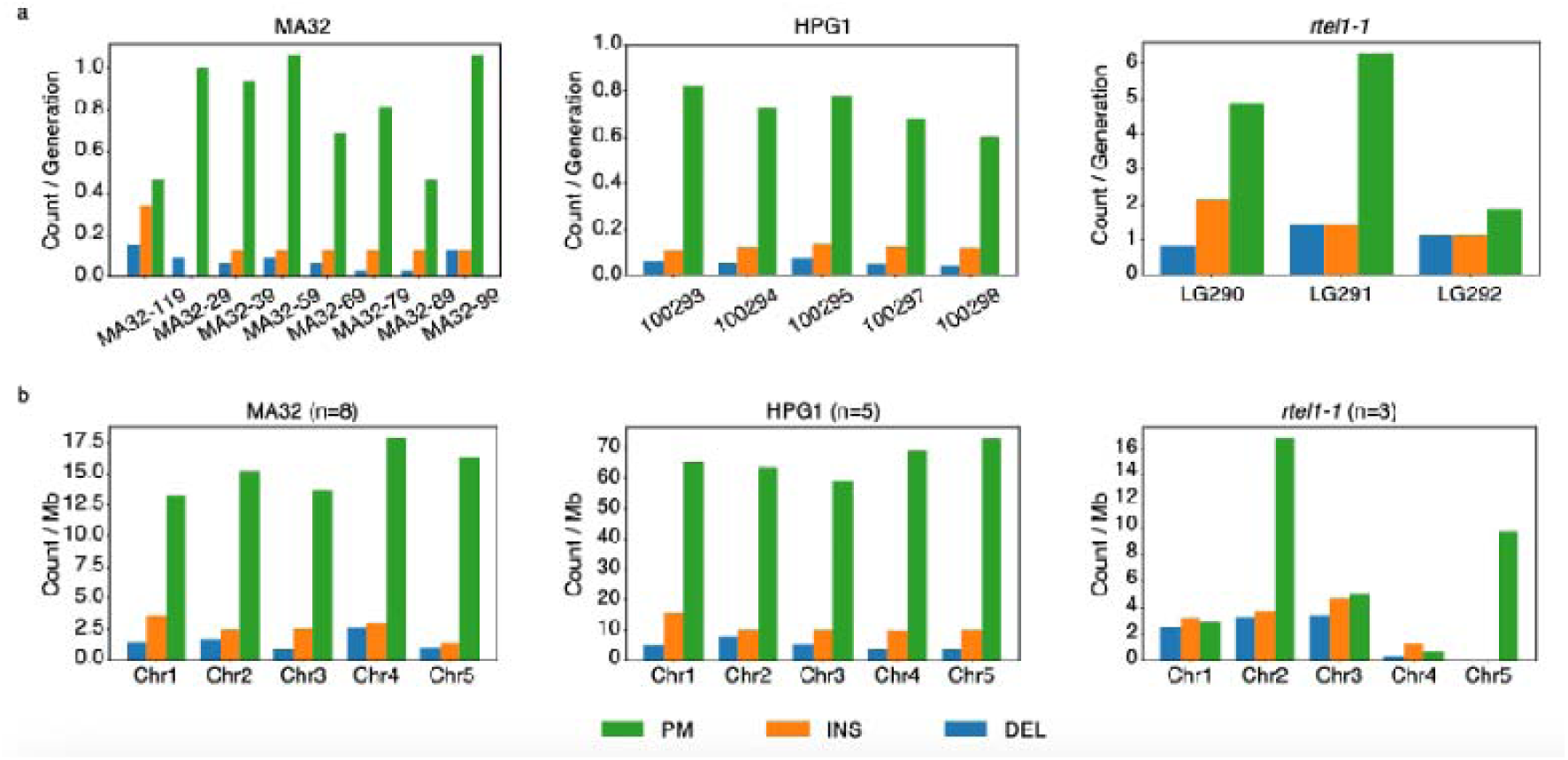
Distribution of true homozygous centromeric mutations across three groups. Mutation counts are shown at the level of individual samples (a) and individual chromosomes (b). For comparability, mutation counts per sample were normalized by the number of selfing generations, whereas mutation counts per chromosome were normalized by centromere size. Mutations are colour-coded by type: point mutations (green), insertions (orange), and deletions (blue). Chi-square goodness-of-fit tests revealed no significant differences in mutation counts among samples or chromosomes for MA32 and HPG1 (P > 0.05), indicating a relatively uniform distribution of centromeric mutations. Statistical power was limited for *rtel1-1* due to the small sample size. PM, point mutation; INS, insertion; DEL, deletion.

**Extended Data Fig. 3.**
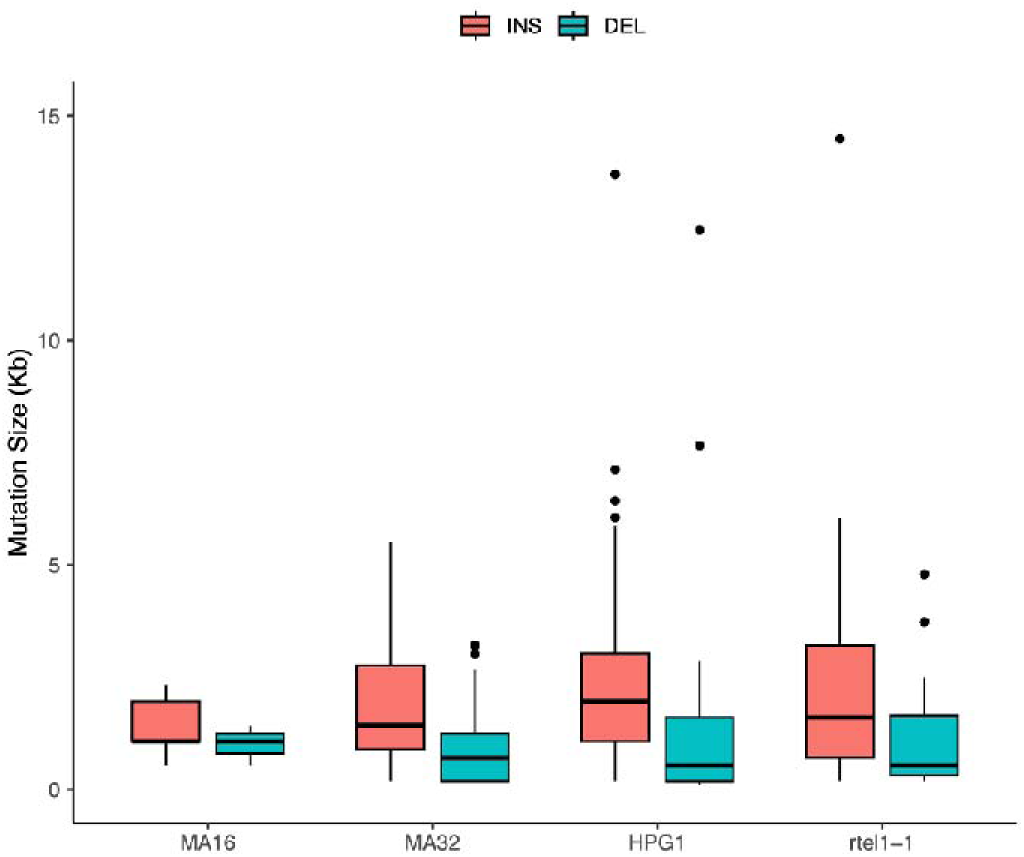
Size distribution of homozygous centromeric indels across four groups. The boxplot illustrates the length of identified indels. Notably, two exceptionally large deletions in *rtel1-1* (59,508 bp and 105,343 bp) are excluded from the plot for clarity.

**Extended Data Fig. 4.**
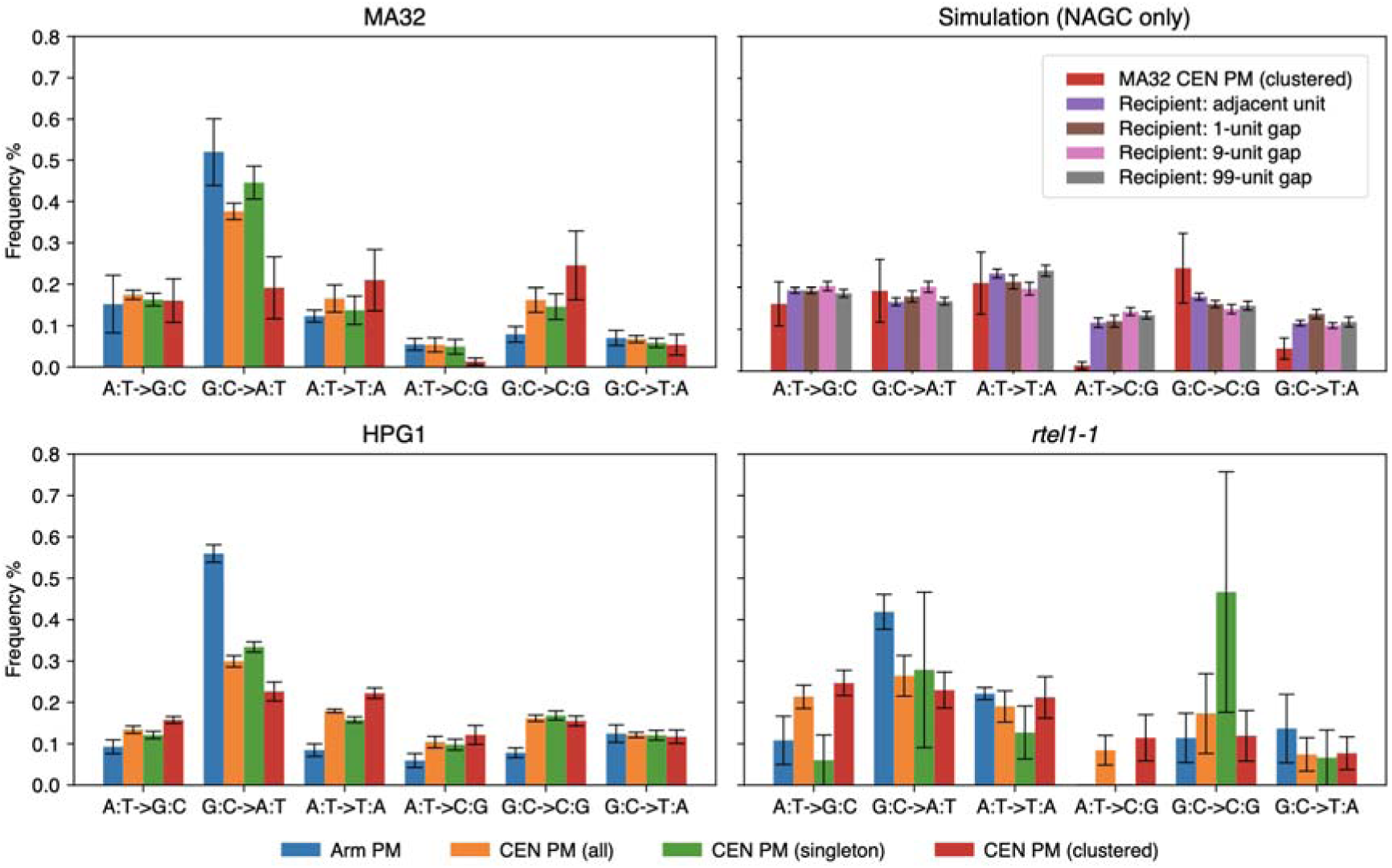
Mutation spectra of homozygous centromeric point mutations across different groups and comparison with NAGC-only simulations. Bar plots show the frequencies of homozygous centromeric point mutations across four datasets (MA32, HPG1, rtel1-1, and simulation). Colours indicate genomic regions or mutation categories: chromosome arms (blue), all centromeric point mutations (orange), singleton centromeric point mutations (green), and clustered centromeric point mutations (red). The upper-right panel shows simulations including NAGC only. Simulated mutation spectra are stratified by recipient unit position relative to the donor, including adjacent units, and units separated by 1, 9, or 99 intervening repeat units. This comparison allows evaluation of whether the mutation patterns generated by NAGC recapitulate the observed clustered mutation spectra.

**Extended Data Fig. 5.**
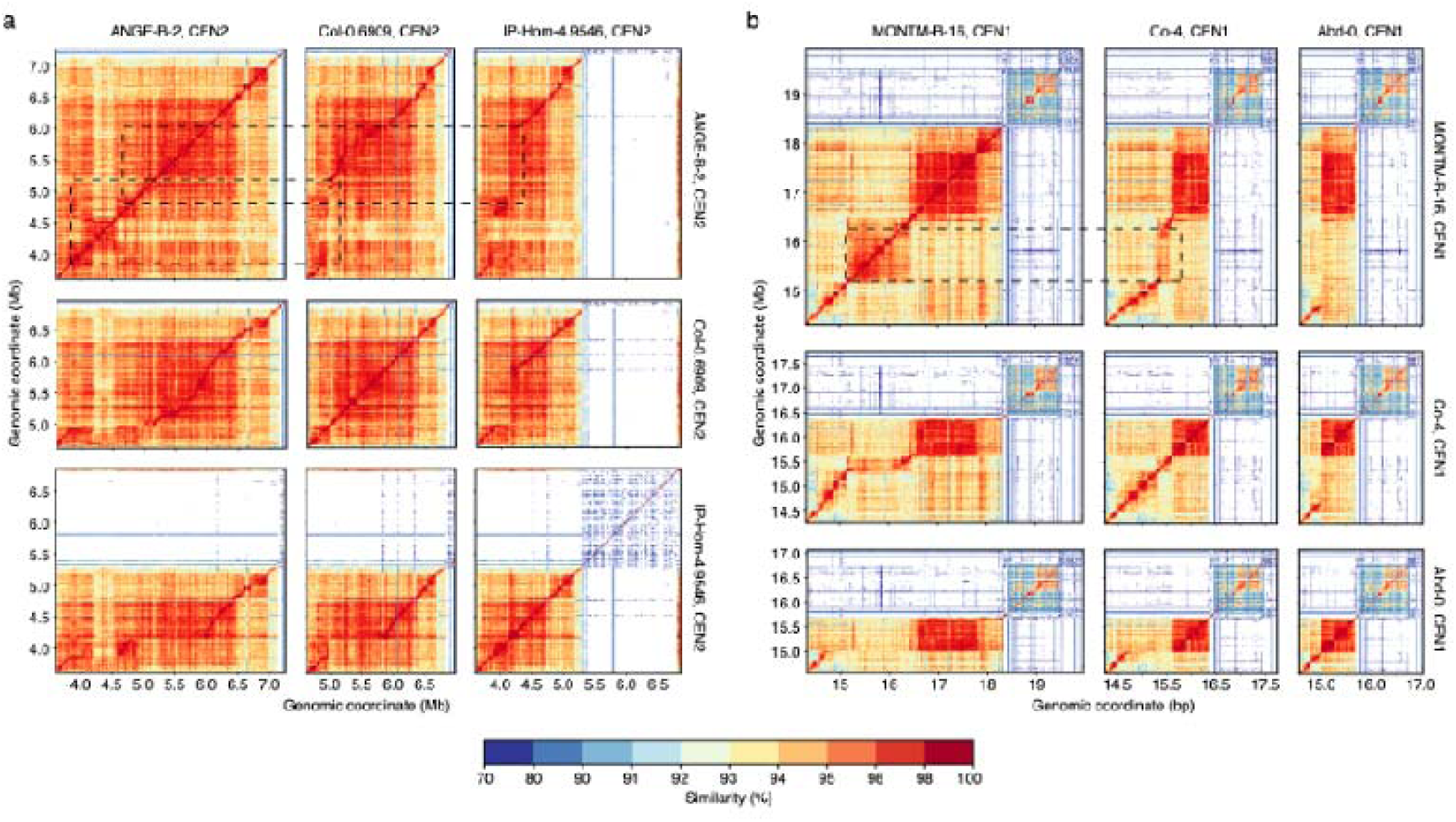
Examples of large rearrangements in centromeric sequences. Heatmaps show self- and pairwise sequence identity for the centromeres of chromosome 2 from ANGE-B-2, Col-0.6909, and IP-Hom-4.9546 (**a**), and for the centromeres of chromosome 4 from MONTM-B-16, Co-4, and Abd-0 (**b**). The putative deletions are highlighted by the dashed boxes. Sequence identity was calculated using ModDotPlot with a 10 kb window size.

**Extended Data Fig. 6.**
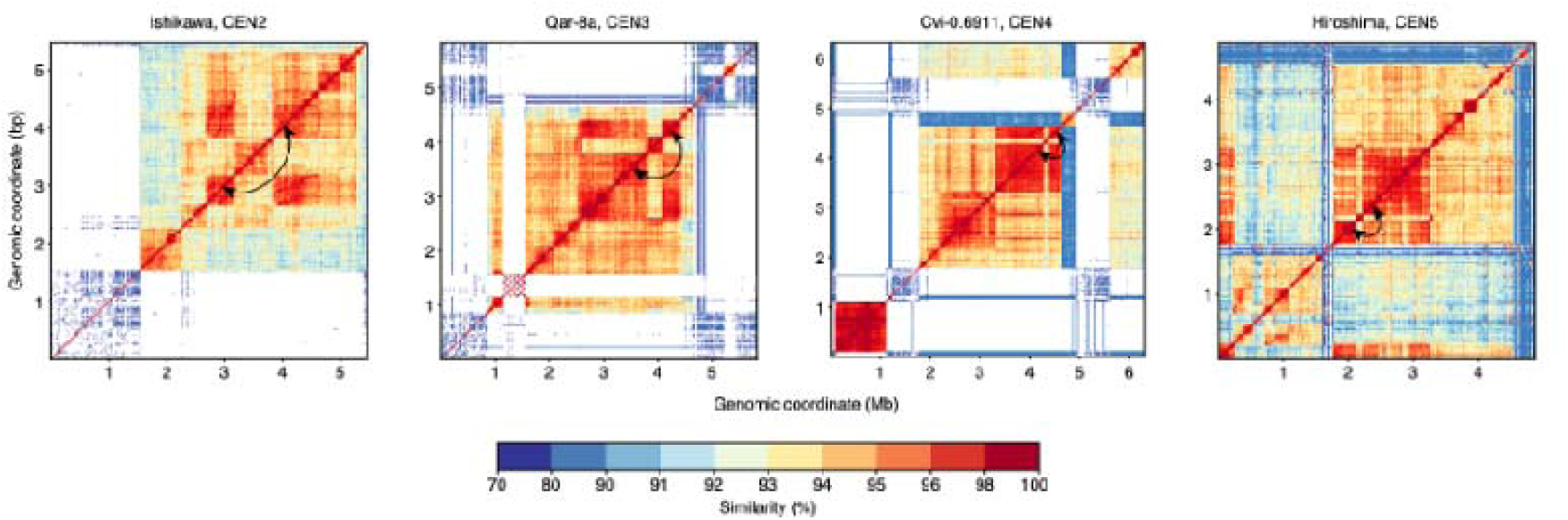
Examples of long-distance similarities in centromeric sequences. Heatmaps show self-sequence identity for the centromeres of chromosome 2 in Ishikawa, chromosome 3 in Qar-8a, chromosome 4 in Cvi-0.6911, and chromosome 4 in Hiroshima. Sequence identity was calculated using ModDotPlot with a 10 kb window size. Regions exhibiting long-distance similarity are indicated by arrows.

**Extended Data Files. Dynamic changes in simulated centromeres.** A total of 15 movies are provided, corresponding to three independent simulations for each of the five centromeric sequences. Files are named according to centromere identity and simulation replicate (CEN1–CEN5; test001–test003), following the format CENX.testY.mp4. Each movie shows the progression of self-sequence identity within a simulated centromeric repeat array over 150,000 generations, starting from the initial sequence. Frames are sampled every 1,000 generations. Each frame represents pairwise sequence identity between non-overlapping 10 kb windows within the array. Sequence identity values are colour-coded according to the scale shown on the right.

