## Supplementary Results for "The mutational dynamics of the Arabidopsis centromeres"

Dong *et al*.

### – Supplementary Information

### Supplementary Results

#### Replicated genome assembly

The sizes of the pseudo-chromosomes were highly consistent between all four assemblies (between 133.6 and 133.9 Mb) (Supplementary Table 2). The small size differences were almost entirely due to differences in the reconstruction of the 45S rDNA tandem arrays (nucleolus organizer regions (NORs)) at the beginning of chromosomes 2 and 4.

After scaffolding, the pseudo-chromosomes included all long contigs of each assembly even though some short contigs were not integrated. Almost all of the unscaffolded contigs showed high sequence similarity to the 45S rDNA sequences, suggesting that the NORs were fragmentarily assembled, but that their high repetitiveness impeded a contiguous assembly, while the rest of the genome was well assembled. This is further supported by the total length of the unscaffolded contigs (~10 Mb in each assembly), which was very similar to the estimated sizes of NOR regions^1^, and the fact that all other highly repetitive tandem arrays were assembled well. For example, all 16 5S rDNA clusters as well as 17 of the 20 centromeres were assembled into single contigs presumably due to shorter repeat units (centromeric repeat units: 178 bp^2–4^; 5S rDNA repeat units: 500 bp^5,6^; 45S rDNA repeat units: >10 kb^7^) and higher unit divergence.

In total, we found 14 to 21 breaks in each of the four assemblies. The regions where the assembly broke were strikingly similar between the four samples (73 breaks in 32 regions, Extended Data Fig. 1a). Around two-thirds of the breaks occurred in regions where other assemblies also broke, including eight regions where all four assemblies broke. Around half of the regions with assembly breaks (15 out of 32) overlapped with GA(A) / CT(T) repeats, which were previously found to impede HiFi sequencing^8,9^, resulting in reduced sequencing depth and in turn in assembly breaks.

Almost all of the errors were indels (96.7%), while single nucleotide errors (SNEs) accounted only for a minor proportion (3.3%). Most of the indel errors were small, but 33 of them were larger than 50 bp, reaching up to 47,397 bp.

Around 77% of the indel errors (75% of all errors) were only 1 or 2 bp in size and occurred in simple repeats, where the simple repeats were assembled either with too many or too few repeat units (Supplementary Fig. 1). Most of these errors resulted from a misrepresentation of the repeat length in HiFi reads (Supplementary Fig. 1b) and most of them were found in homopolymers or dinucleotide repeats but also in tri- or seven-nucleotide repeats (Supplementary Fig. 1). Errors in simple repeats were already observed when they were only three units long, but the error rate increased when the number of repeat units was above 20 (Supplementary Fig. 2). Around 20% of the errors in simple repeats were identified in regions where one or even two of the other assemblies were incorrect as well, implying that simple repeats with high repeat number are extremely error-prone.

Unexpectedly, the ratio of insertion *vs.* deletion errors varied between the replicates. For example, in A2, 90% of the indel errors were insertion errors, while in A1 only 23% of the indel errors were insertions. The ratio of indel errors, however, mirrored the ratio of indel errors in the reads, suggesting that the rate of indel errors is a direct consequence of read quality (Supplementary Fig. 3).

Many of the errors were close to assembly gaps. Assembly sequence close to gaps is usually generated from fewer reads than the rest of the assembly and errors in these reads are more likely to be introduced into the assembly. Across the four assemblies, we identified 268 errors (17.5% of all errors) close to gaps (<2.5 kb, Supplementary Fig. 4). Notably, the majority of the SNEs (70.6%) were introduced in these regions.

Only a very small number of assembly errors resulted from the assembly process itself (3.4%, 52 out of 1,532). Most of them (31 out of 52) were small-scale, where the sequence of the raw reads did not match the assembled sequence (Supplementary Fig. 5). Other errors resulted from mistakes in assembly graph traversal or scaffolding together, causing many of the few large-scale errors (>50 bp) (Supplementary Fig. 6, Supplementary Fig. 7).

Finally, correcting the errors with conventional polishing approaches (based on the alignments of short and/or long reads) identified less than a quarter of all errors and at the same time “over-corrected” multiple hundreds of positions, leading to more additional than corrected errors (Supplementary Fig.8).

#### Causes of errors in the HiFi genome assemblies

Across the four assemblies analysed, we identified 1532 errors, the vast majority of which were small-scale (1-2 bp). Upon investigation, these errors fell into three categories: (1) copy number errors in simple repeats, (2) sequencing errors retained in low-coverage regions, and (3) assembly tool errors and reference-based scaffolding errors.

##### 1. Simple repeats introduce copy number errors

Simple repeats, such as homopolymers and di- or trinucleotide repeats, were particularly prone to sequencing errors, which appeared as copy number discrepancies — reads over- or underestimating the actual repeat length, leading to inconsistent consensus sequences. This was the most frequent error type, accounting for 198 (B1, 74.1%) to 466 (A2, 81.6%) errors, over 75% of which were 1-2 bp in size, with a maximum of 21 bp. Errors followed the monomer pattern length, predominantly occurring in homopolymer and dinucleotide repeats, while trinucleotide repeats were less affected. Copy number errors increased when repeat counts exceeded ten copies.

Interestingly, the lower error rate in centromeres correlated with a notable reduction in simple repeats compared to chromosome arms. Apart from peri-centromeric regions, true centromeres contained virtually no simple repeats (Extended Data Fig. 1a).

##### 2. Low sequencing depth preserves random errors

PacBio HiFi reads typically maintain an error rate of ~0.1% (<https://www.pacb.com/blog/long-read-sequencing-myths-debunked-part-1-hifi-sequencing/>), correctable during consensus assembly. However, HiFi reads often drop in GA(A)/CT(T) repeat regions^8,9^, leading to reduced depth and assembly gaps. These regions were prone to retaining occasional sequencing errors, as few supporting reads remained to correct them. Across the four assemblies, we identified 33-96 such errors, accounting for 11.3%-24.6% of total errors. Most occurred within 25 kb of GA(A) repeats and were backed by fewer than ten reads.

##### 3. Assembly and scaffolding introduce a few large errors

In addition to sequencing-related errors, assembly tools themselves introduced a small fraction of errors across our datasets. We identified 31 assembly errors across all four replicates, the majority (67.74%) being small 1-2 bp errors, though one exceptional case reached 1,491 bp. These errors appeared consistently in both unitigs and contigs generated by hifiasm^10^ but were not supported by HiFi reads (Supplementary Fig. 5). The root cause of these errors remains unclear, but their overall frequency was extremely low, and they were readily detectable by aligning the HiFi reads back to the assembly.

We also observed errors near branching points ("forks") in the assembly graph. Repeats created ambiguous paths, resulting in either expansion or collapse errors during graph traversal. We detected four collapse errors, one base substitution, and a 36 kb expansion error. An example of a 5,980 bp error in the 5S rDNA region on chromosome 4 (B1 assembly) showed that complex local patterns misled the assembly graph, forming a "bubble" that caused a major collapse (Supplementary Fig. 6). By analysing unitig alignments to the F0-16 and B1 assemblies, we determined that misidentification of parallel paths led to the bypassing of a key unitig, creating the collapse.

Additionally, reference-based scaffolding — used to build chromosome-level assemblies in this study — introduced five misplacements or misorientations among contigs (22 to 47 kb in length) and 11 redundant overlaps between contigs (1 to 32 kb in length). This approach, which aligns contigs to a known reference genome, can fail when the target genome diverges from the reference, especially for smaller contigs. While scaffolding errors were rare, they highlight the limitations of relying on reference-based methods for highly repetitive regions.

#### Error correction tools introduced more errors than they corrected

We evaluated several widely used error-correction and polishing tools — Pilon^11^, DeepVariant^12^, pbsv (<https://github.com/PacificBiosciences/pbsv>), and Sniffles^13^ — by mapping Illumina short reads or HiFi long reads to assemblies and identifying consistent discrepancies. Among these, Pilon^11^ showed the highest sensitivity, detecting about one-fourth of the errors. However, it also introduced ~85% more errors through overcorrection (Supplementary Fig. 8). Pbsv and Sniffles^13^, designed for structural variant detection from long reads, identified five indel errors (11-50 bp) across four replicates but showed limited performance in correcting other types of errors.

This analysis revealed that mismatches between reads and assembly — likely stemming from the assembly algorithm — were the most detectable, with 88% recognized. Errors at assembly gaps with low depth were partially identified (20-40%), while errors arising from simple repeats were detected in less than 20% of cases. No tool successfully identified large errors (≥ 50 bp) caused by assembly graph traversal or scaffolding.

Further investigation of Pilon’s^11^ overcorrected regions showed that most had low mappability, despite HiFi reads aligning well. These results suggest that current tools struggle to resolve errors in repetitive regions and large indels, often risking overcorrection and introducing additional errors.

### Supplementary Methods

#### Addressing repetitive and structurally complex regions

Highly repetitive and structurally complex regions, such as centromeres, posed challenges for reliable sequence alignment. These regions often produced clusters of false positive mutations or assembly errors. To resolve this, we extracted sequences from flagged error/mutation clusters and realigned them to the F0 genome using minimap2^14^. By shortening the mapped regions and re-aligning, we identified larger, genuine mutations or assembly errors that better explained the observed sequence differences.

Moreover, regions with consistently incomplete assembly — including nucleolus organizer regions (NORs) and variable-length telomeres — were excluded from error and mutation counts. This exclusion covered the first 500 Kb of Chr2, the first 100 Kb of Chr4, and the telomeric ends of the remaining eight chromosome arms.

#### Sequencing depth calculation for samples A and B

To measure sequencing depth for samples A and B, we generated base-by-base coverage statistics from BAM files produced by aligning HiFi and Illumina reads to the ancestral genome assemblies. We used bedtools^15^ v2.29.0’s depth function with the “-aa” parameter to include all positions. Additionally, we calculated the average sequencing depth within 1 Kb non-overlapping windows using bedtools^15^ v2.29.0’s map function.

#### Indel biases analysis in HiFi read alignment for samples A and B

To account for differences in insertion and deletion error biases between samples A and B such as the higher frequency of deletion errors in A1 and insertion errors in A2 we quantified indels in the HiFi read alignments using samtools^16^ v1.19.2’s mpileup function. A custom Python script was then used to analyze indel sizes within the reads. We identified indel sites by counting positions with ≥10 aligned reads, where >50% of reads displayed an indel at that position, ensuring robust detection of alignment mismatches.

#### Error correction with existing tools

We evaluated the performance of several existing assembly error correction tools, which generally operate by aligning sequencing data to the assembly and flagging mismatches as potential errors. The tested tools included Pilon^11^ v1.24 (--fix snps, indels --vcf --changes --tracks), which relies on short reads; pbsv v1.16 (<https://github.com/PacificBiosciences/pbsv>) and Sniffles^13^ v2.0.7, both designed for long-read data; and DeepVariant^12^, using --model_type WGS, PACBIO, or HYBRID_PACBIO_ILLUMINA for short-read, long-read, and hybrid data, respectively. The errors identified or corrected by these tools were compared against a manually curated list derived from assembly comparisons between replicates to assess accuracy and reliability.

### Supplementary Figures

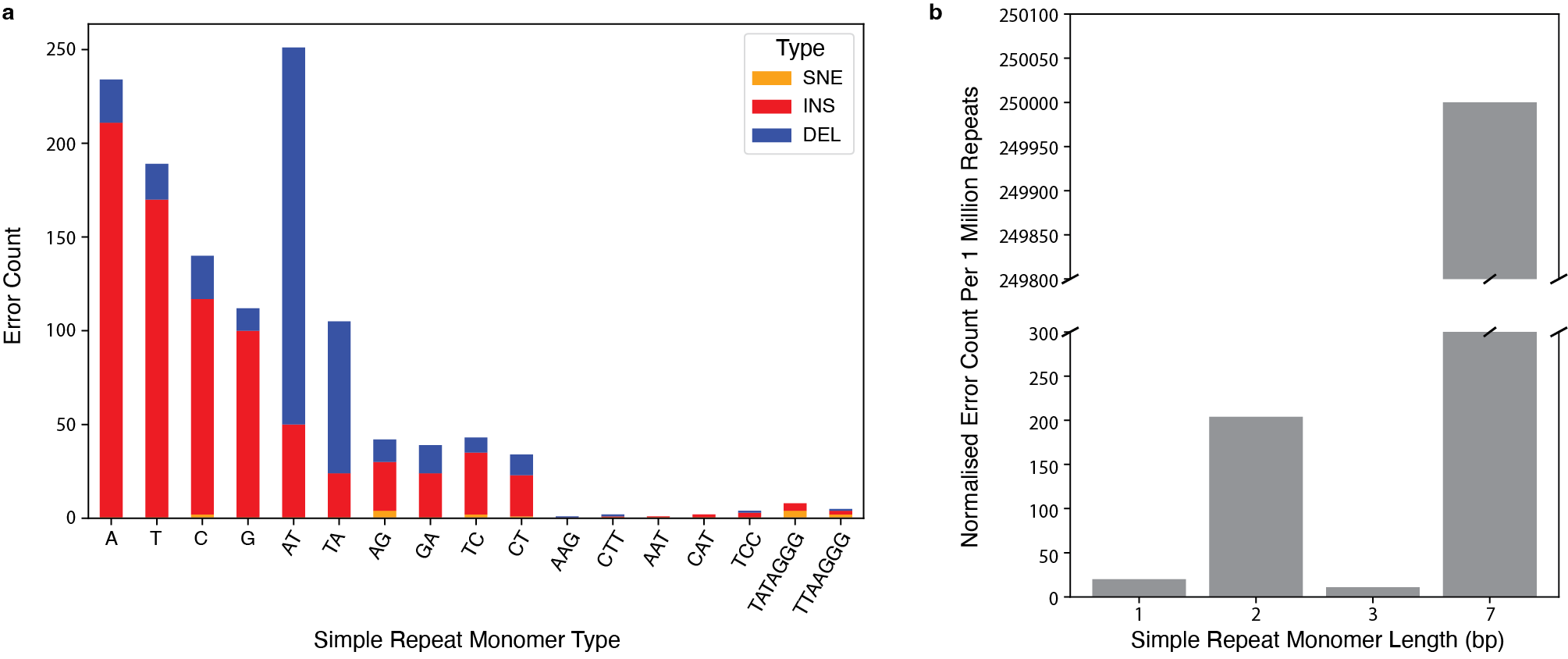

**Supplementary Fig. 1. Most HiFi-based assembly errors are found in homopolymers or dinucleotide repeats.** **a.** Bar plots show the error count from each type of repeat monomer, with error types color-coded: SNE (orange), insertion error (red), and deletion error (blue). **b.** Bar plots show the normalized number of errors from different monomer lengths along the chromosome. Error numbers are normalized to the number of errors per one million repeats to account for the variation in simple repeat types across the chromosome.

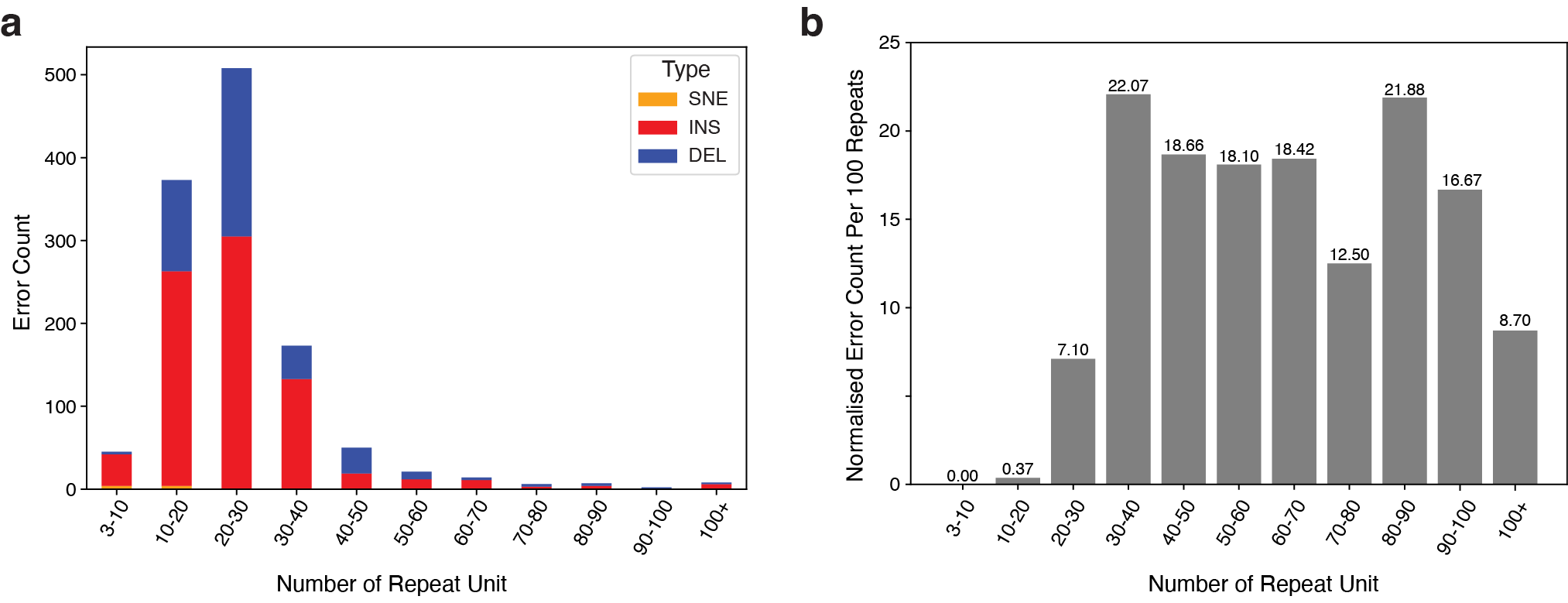

**Supplementary Fig. 2. HiFi-based assembly errors in simple sequence repeats are more frequent in longer repeat units.** **a.** Bar plots show the number of errors in different numbers of repeat units, with error types color-coded: SNE (orange), insertion error (red), and deletion error (blue). **b.** Bar plots display the normalized error count from each group of repeat unit numbers.

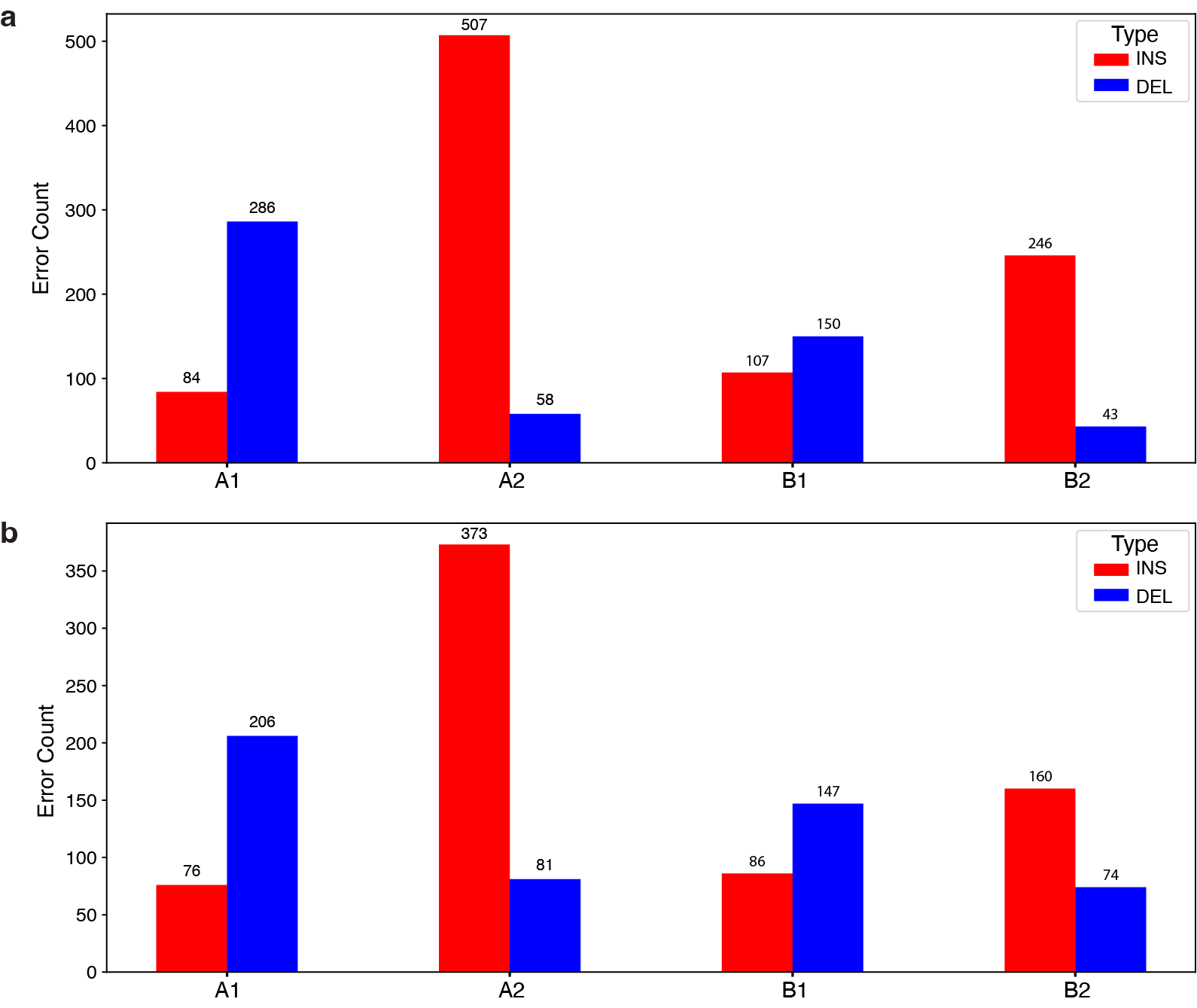

**Supplementary Fig. 3. HiFi-based assembly indel errors mirror the indel errors in the reads.** **a.** Bar plots show the number of indels in four samples, with error types color-coded: insertion error (red) and deletion error (blue). **b.** Bar plots show the number of positions with more than 10x coverage where more than half of the reads contain either an insertion or deletion error in the alignment.

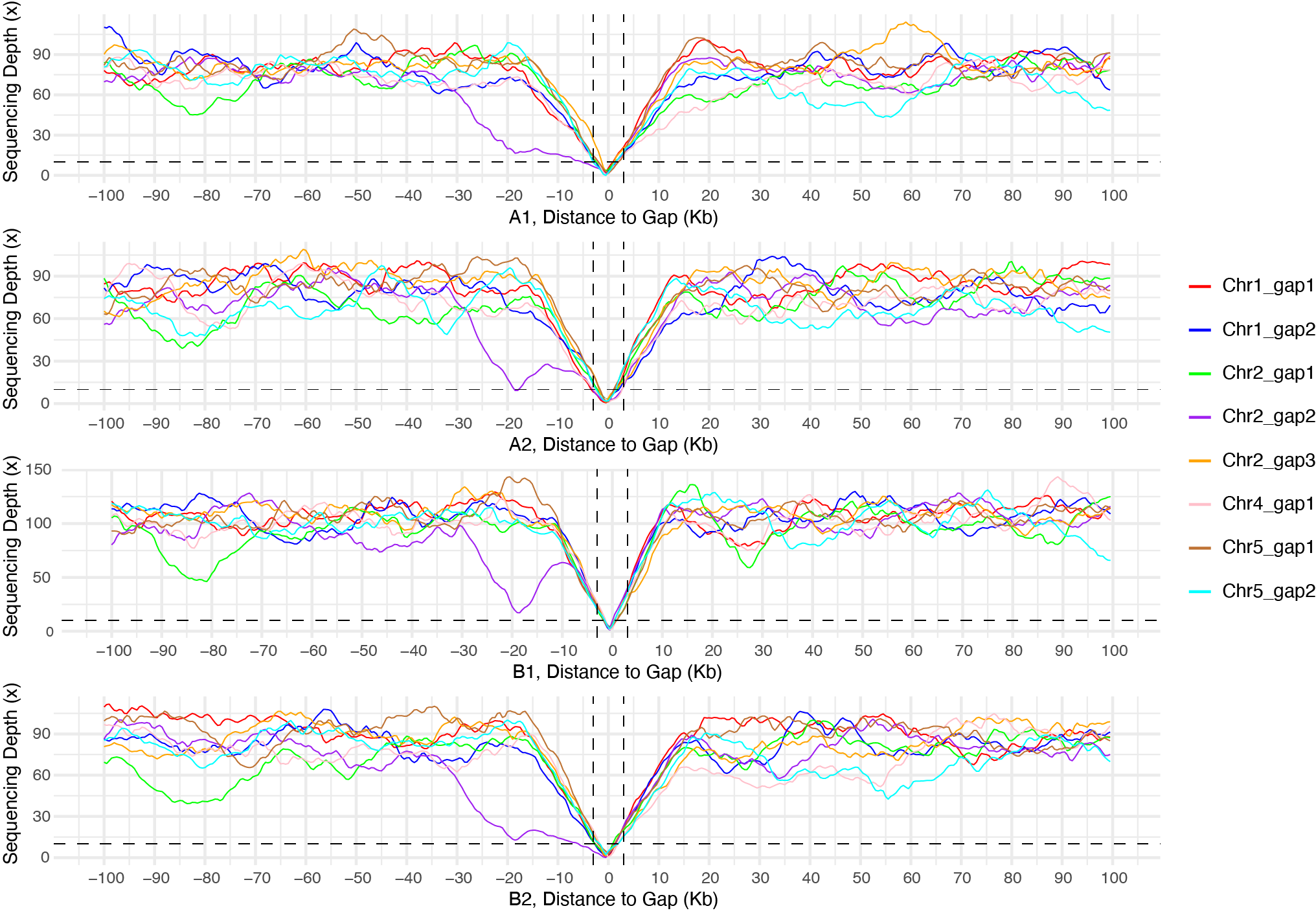

**Supplementary Fig. 4. Sequencing depth of the four samples versus distance from the HiFi-based assembly gap induced by GA repeat.** The x-axis corresponds to the flanking distance from the assembled gap, and the y-axis represents the average HiFi sequencing depth within each 1 kb window. In addition to the low depth (<10x) within the 2.5 kb region flanking the gap, several high GA repeats (e.g., Chr2_gap1_-80kb and Chr2_gap2_-18kb) also show a decrease in sequencing depth.

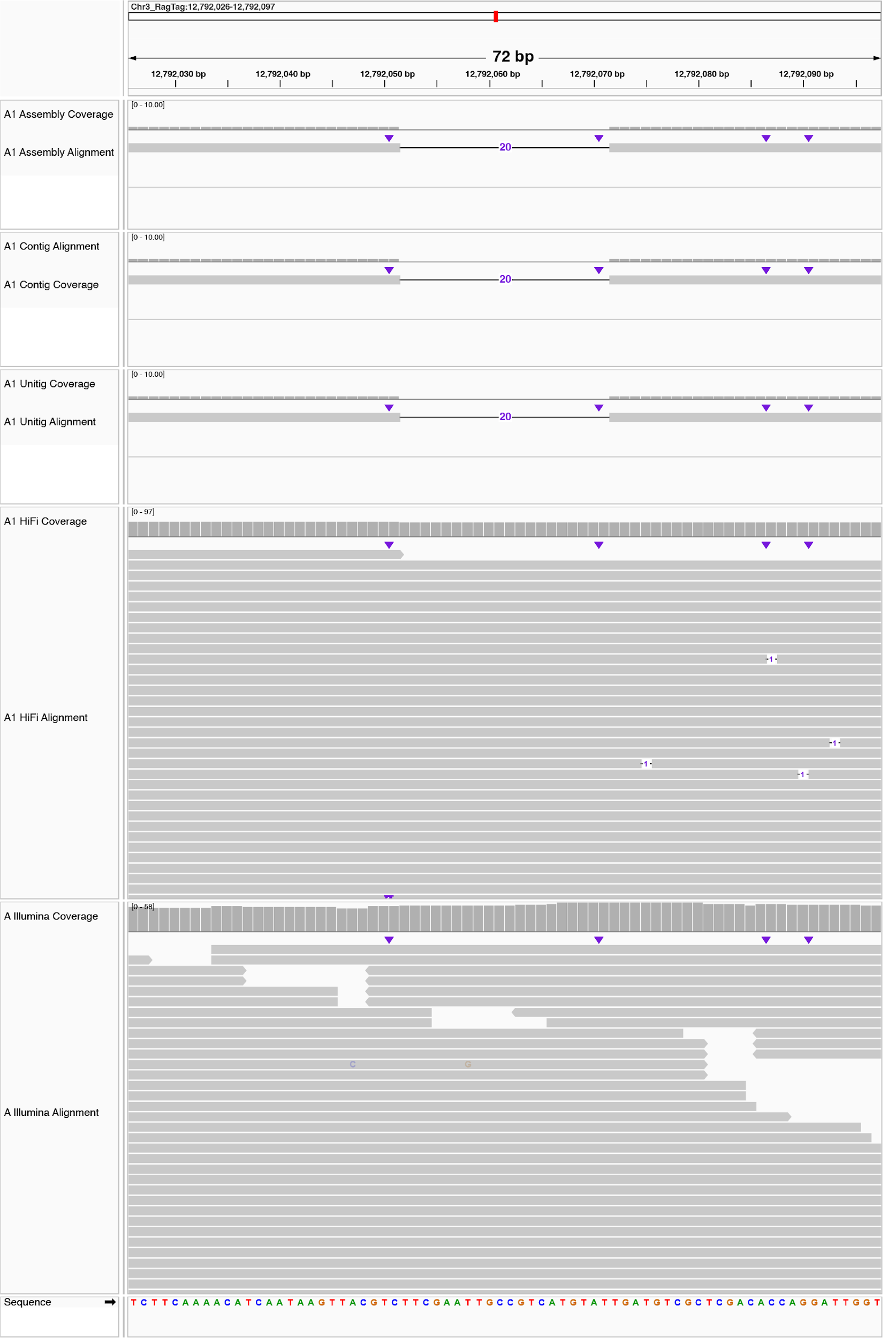

**Supplementary Fig. 5. Examples of HiFi-based assembly errors introduced by the assembly algorithm.** This figure shows a 20 bp deletion error corresponding to position 12.8 Mb on Chromosome 3. The alignment of the A1 chromosome version assembly, the contig-level assembly, the unitig-level assembly, HiFi reads, and Illumina reads are shown from top to bottom. The assembly error is present in the unitig, but there is no HiFi or Illumina read support for the misassembly.

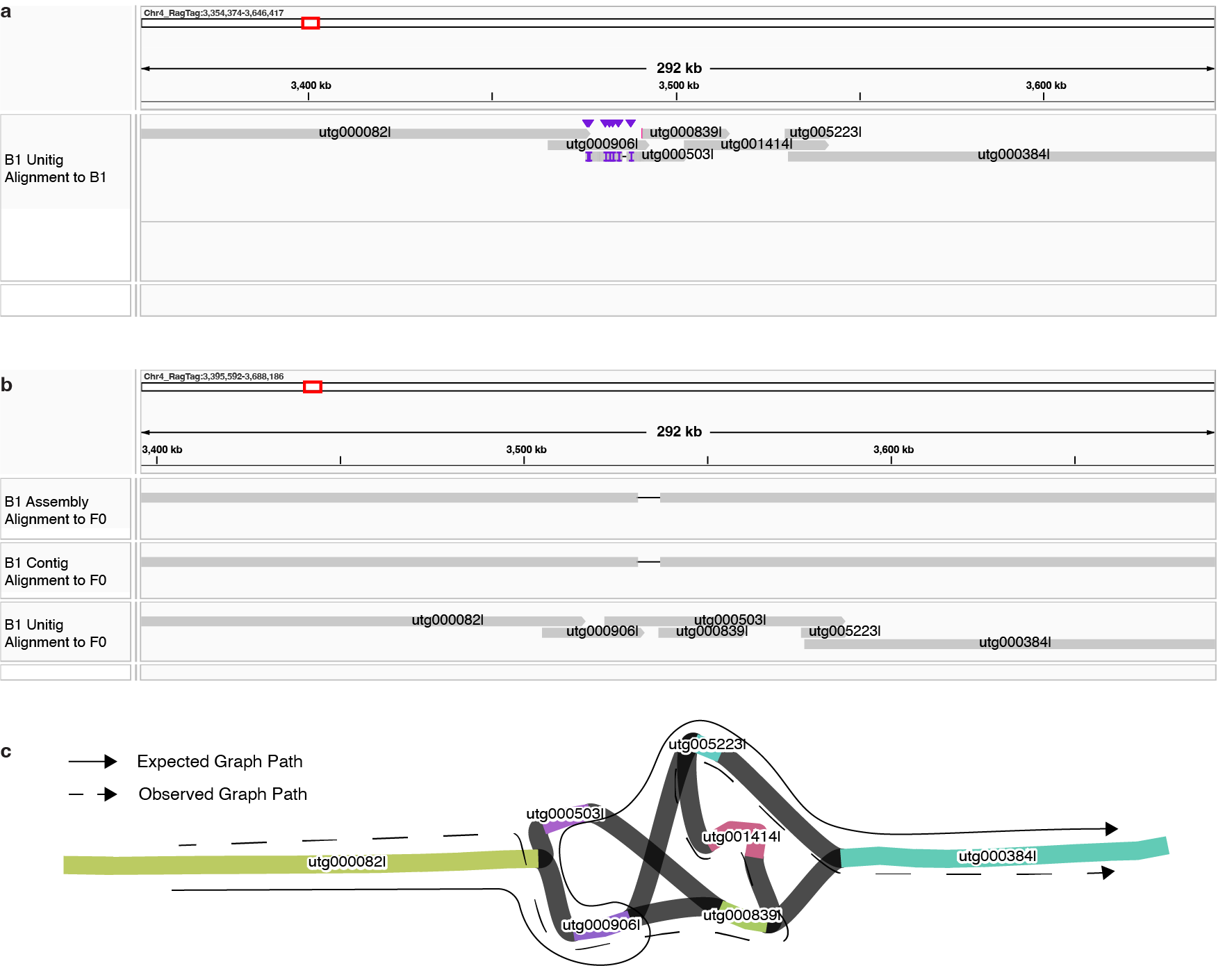

**Supplementary Fig. 6. Example of a HiFi-based assembly error caused by assembly graph.** **a**. Alignment of B1 unitigs to the B1 assembly in IGV reveals the graph path used to generate contig. **b**. Alignment of B1 assembly, B1 contigs and B1 unitigs to the ancestor genome (F0) in IGV (top to bottom), highlighting a 5,980 bp deletion error in both the B1 assembly and contig. **c**. Visualization of the assembly graph in Bandage, showing the expected graph path (solid arrow) and observed graph path (dashed arrow), as inferred from the alignment in panels a and b.

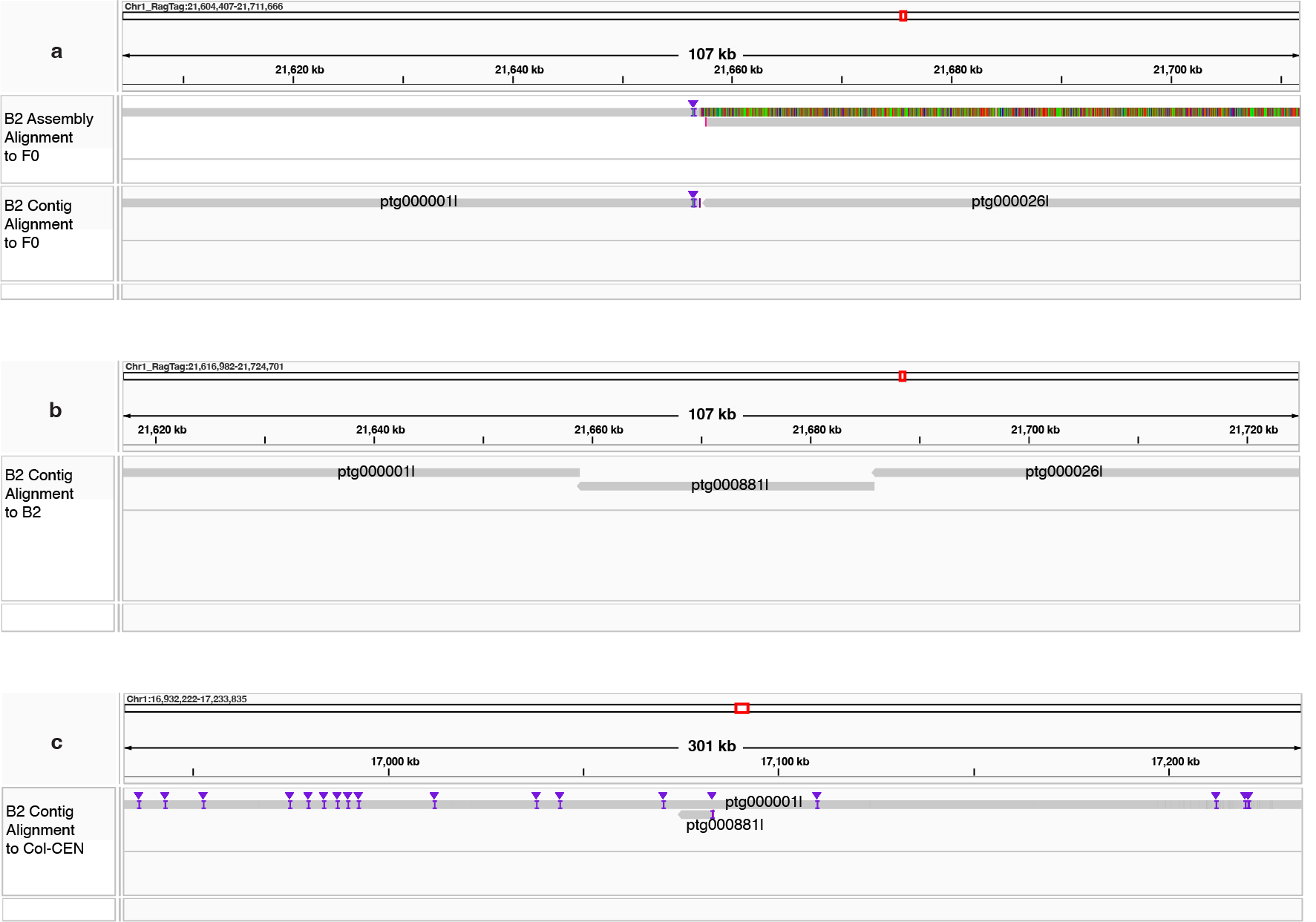

**Supplementary Fig. 7. Example of HiFi-based assembly errors introduced by reference-based scaffolding. a.** Alignment of the B2 assembly and B2 contigs to the ancestor genome (F0) in IGV (top to bottom), showing a large unmapped segment that was soft-clipped. **b.** Alignment of B2 contigs to the B2 assembly reveals that three contigs (ptg000001l, ptg000881l and ptg000026l) were scaffolded together. c. Alignment of B2 contigs to the Col-CEN reference genome (used for scaffolding) shows that ptg000881l and ptg000001l overlap, yet they were scaffolded, resulting in redundant sequence in the assembly.

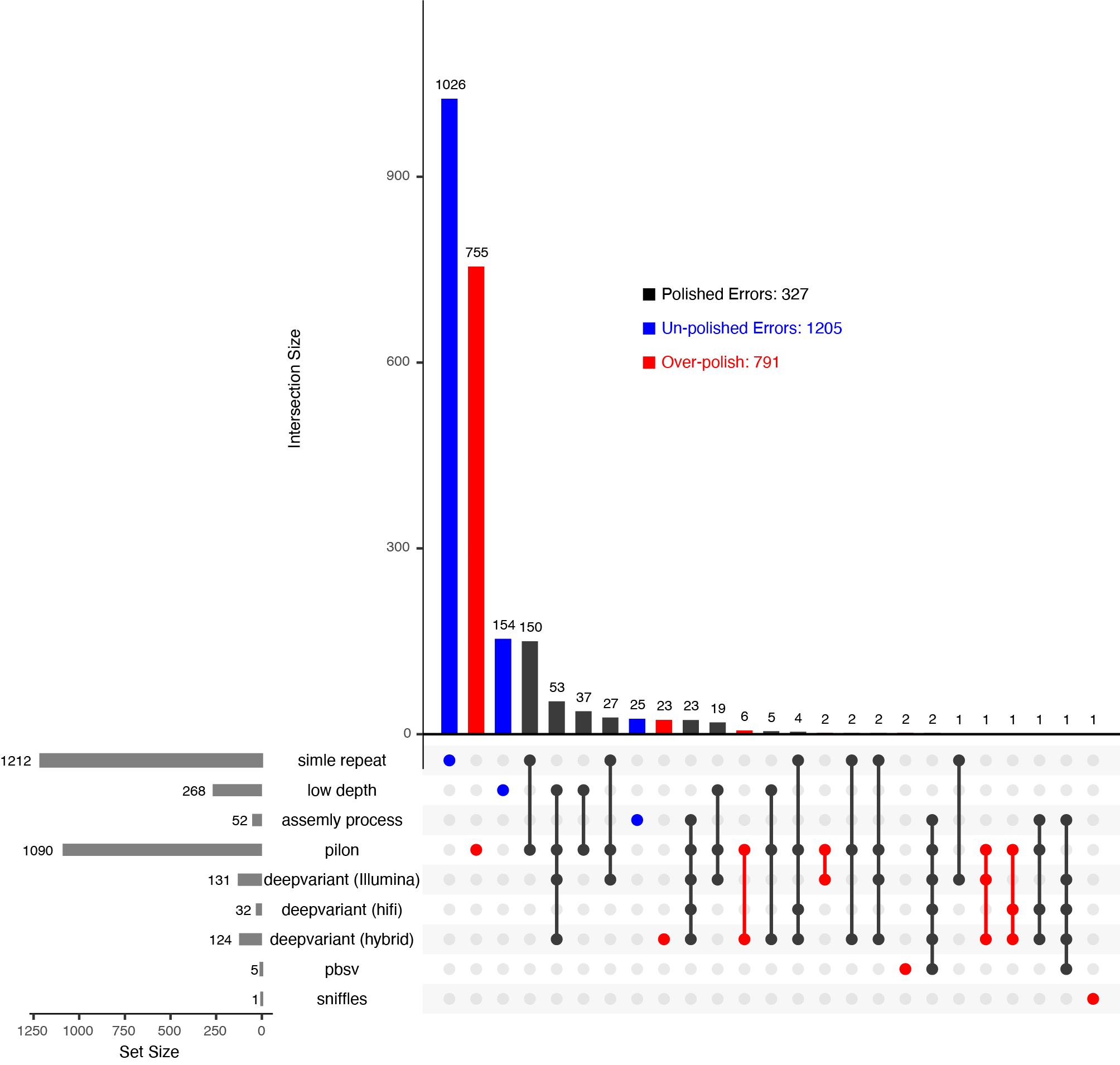

**Supplementary Fig. 8. Existing tools introduced more errors than they corrected in HiFi-based assemblies.** UpSet plot comparing known errors from three sources to errors detected by existing tools. Each intersection represents errors shared between one or more sources and tools, while unique errors from each source or tool appear as single points. The bar plot above quantifies errors per combination, showing that many known errors remained uncorrected, while a substantial portion were mis-corrected, introducing additional errors.

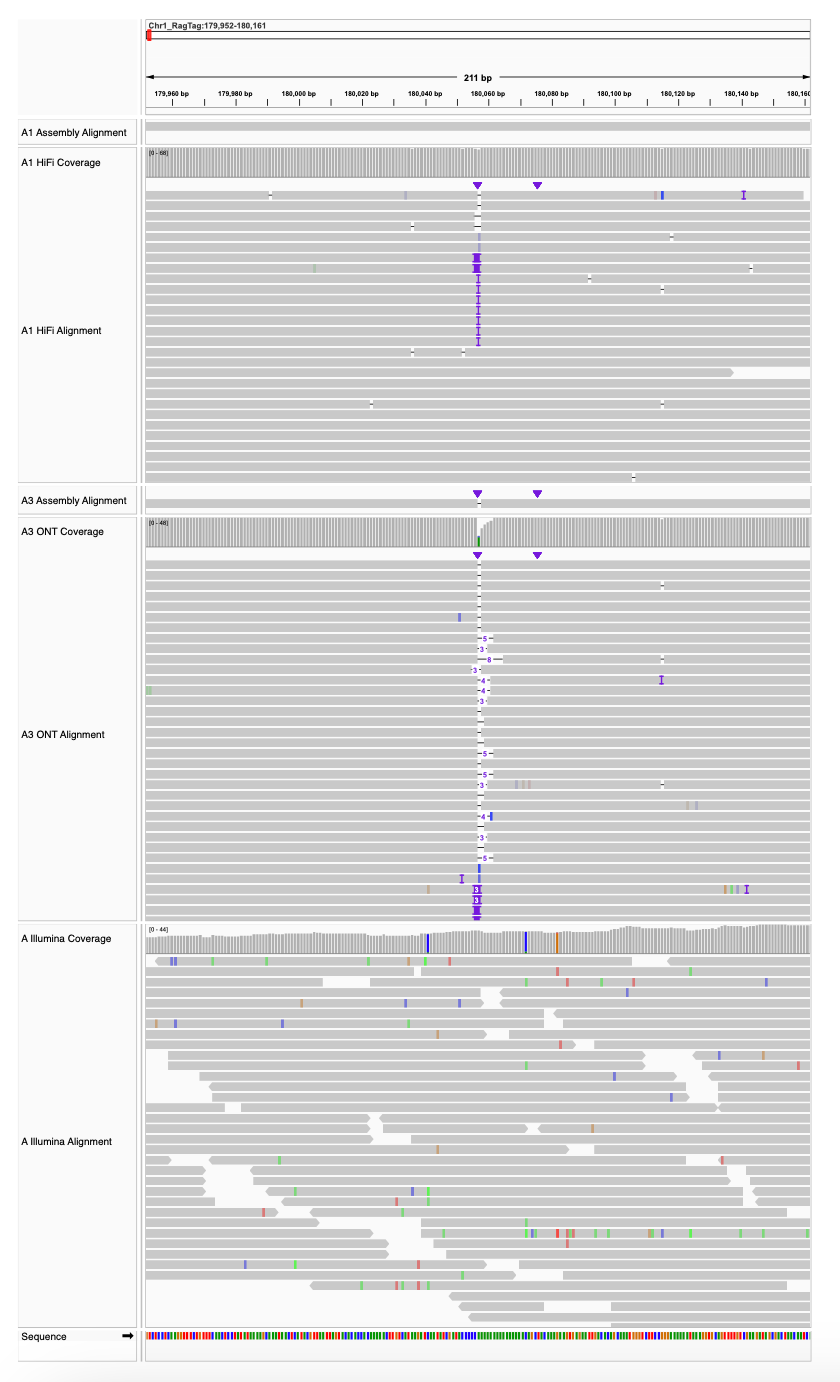

**Supplementary Fig. 9. Examples of ONT-based assembly errors at simple sequence repeats.** This figure shows an example of a homopolymer region consisting of 15 consecutive adenines. The alignments of the HiFi-based assembly (A1), corresponding HiFi reads, ONT-based assembly (A3), and ONT and Illumina reads are shown from top to bottom. Compared to the HiFi reads, ONT reads exhibit reduced and variable copy numbers in the homopolymer tract, leading to inaccurate representation of repeat length and resulting in assembly errors in the ONT-based assembly.

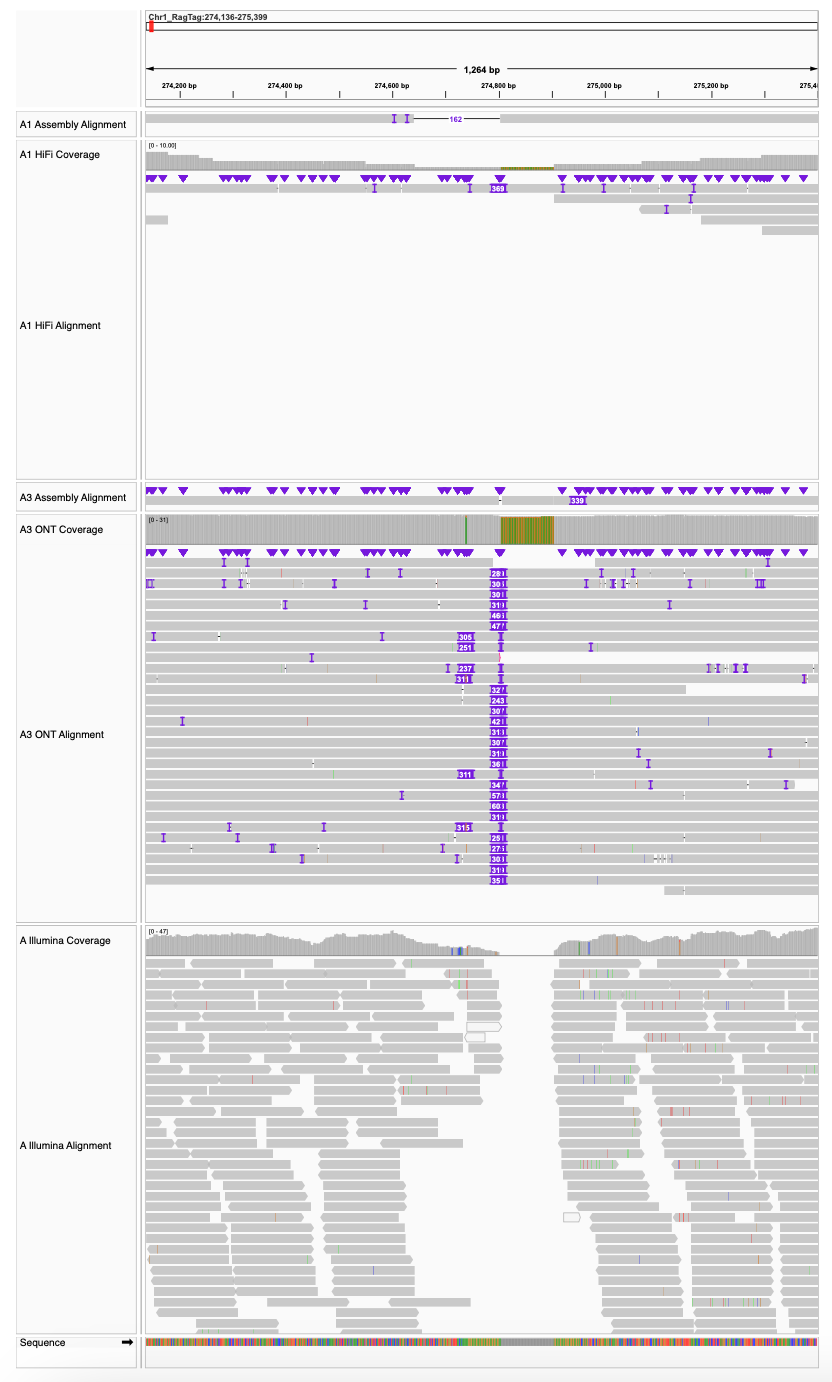

**Supplementary Fig. 10. Example of ONT-based assembly in a GA repeat region.** This figure shows an example of a GA dinucleotide repeat region. The alignments of the HiFi-based assembly (A1), corresponding HiFi reads, ONT-based assembly (A3), and ONT and Illumina reads are shown from top to bottom. While ONT reads provide continuous coverage across the region, substantial variation in repeat copy number is observed among reads, making it difficult to resolve the exact repeat length and leading to ambiguity in the ONT-based assembly compared to the HiFi assembly.

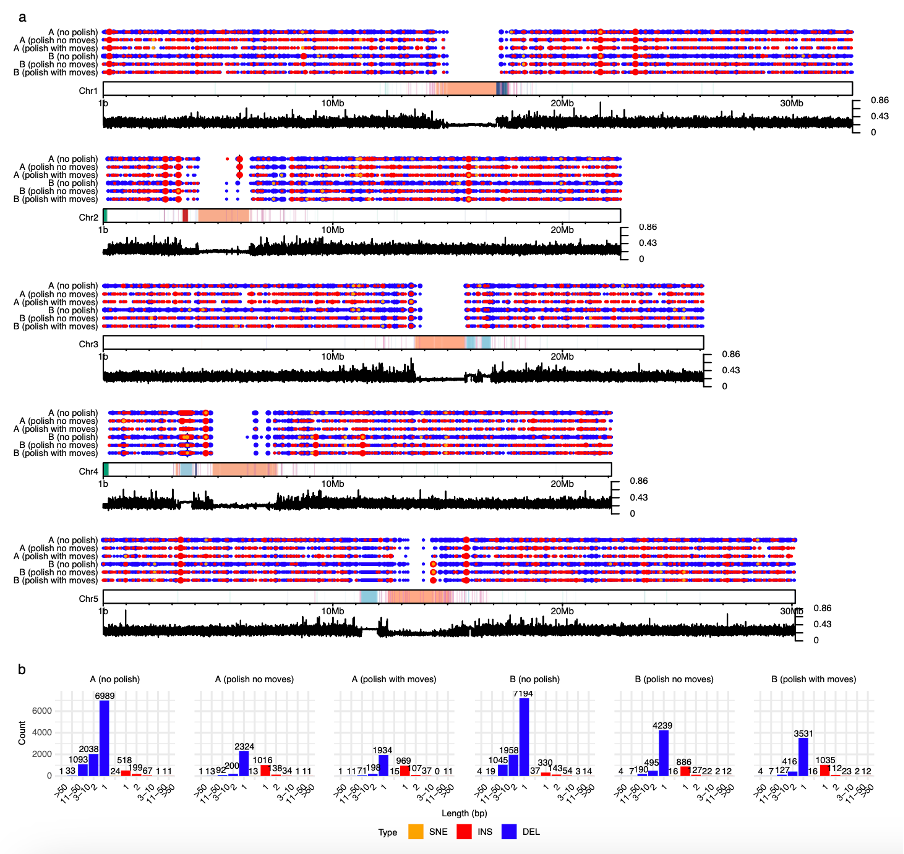

**Supplementary Fig. 11. Assembly errors in ONT-based genome assemblies.** **a.** Distribution of assembly errors across the five chromosomes. Tracks correspond to two assemblies and four polished versions, including polishing without and with move table information (“no moves” and “with moves”, where “moves” refers to move table information generated by Dorado using the --emit-moves option). Each circle represents an assembly error, colour-coded by type: single-nucleotide errors (yellow), insertions (red), and deletions (blue). Circle size reflects error size. Vertical black lines indicate assembly gaps. The central tracks show chromosomal coordinates with repetitive regions highlighted: centromeres (peach), intact Athila elements (magenta), 5S rDNA (light blue), 45S rDNA (green), interstitial telomeric sequences (dark blue), and mitochondrial insertions (red). The line plots below show the proportion of simple sequence repeats within 10 kb windows. **b.** Bar plots showing the distribution of assembly errors of different sizes before and after polishing in the two samples. CEN, centromere; ITSs, interstitial telomeric sequences; MT, mitochondrial sequences; SNE, single-nucleotide error; INS, insertion; DEL, deletion.

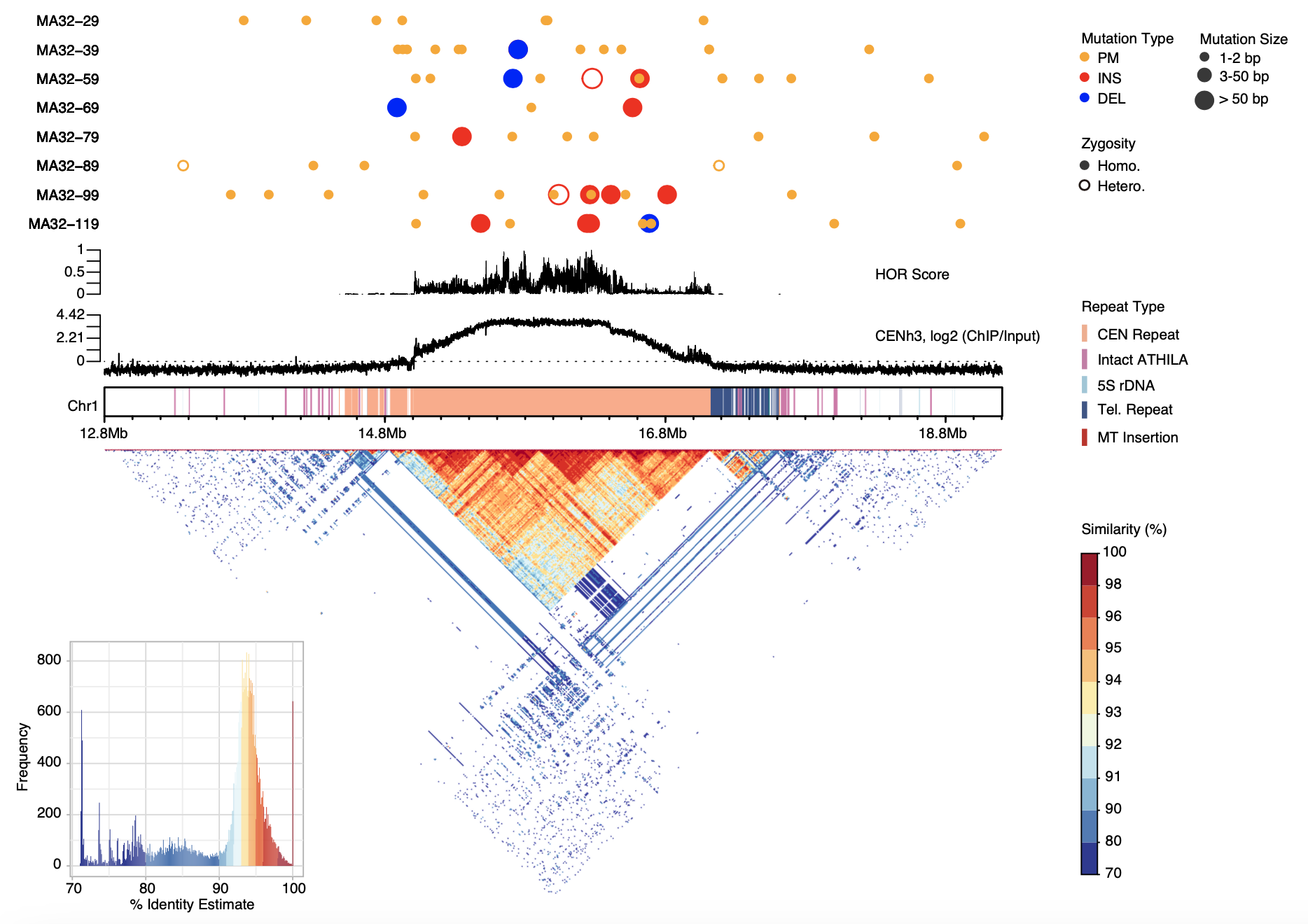

**Supplementary Fig. 12. Mutations within centromere 1.** Eight rows of circles display mutation patterns in the centromere of chromosome 1 in MA32 samples, colour-coded as point mutations (PMs; orange), insertions (red), and deletions (blue). Circle size reflects mutation length: small (1-2 bp), medium (3-50 bp), and large (>50 bp). Solid circles indicate homozygous mutations, while hollow circles indicate heterozygous ones. The two line plots below show higher-order repeat (HOR) scores and log2 CENH3 ChIP-seq enrichment, respectively. Rectangles highlight repetitive regions, and the heatmap below represents pairwise sequence identity between non-overlapping 10 kb regions, with a histogram summarizing identity values in the lower-left corner.

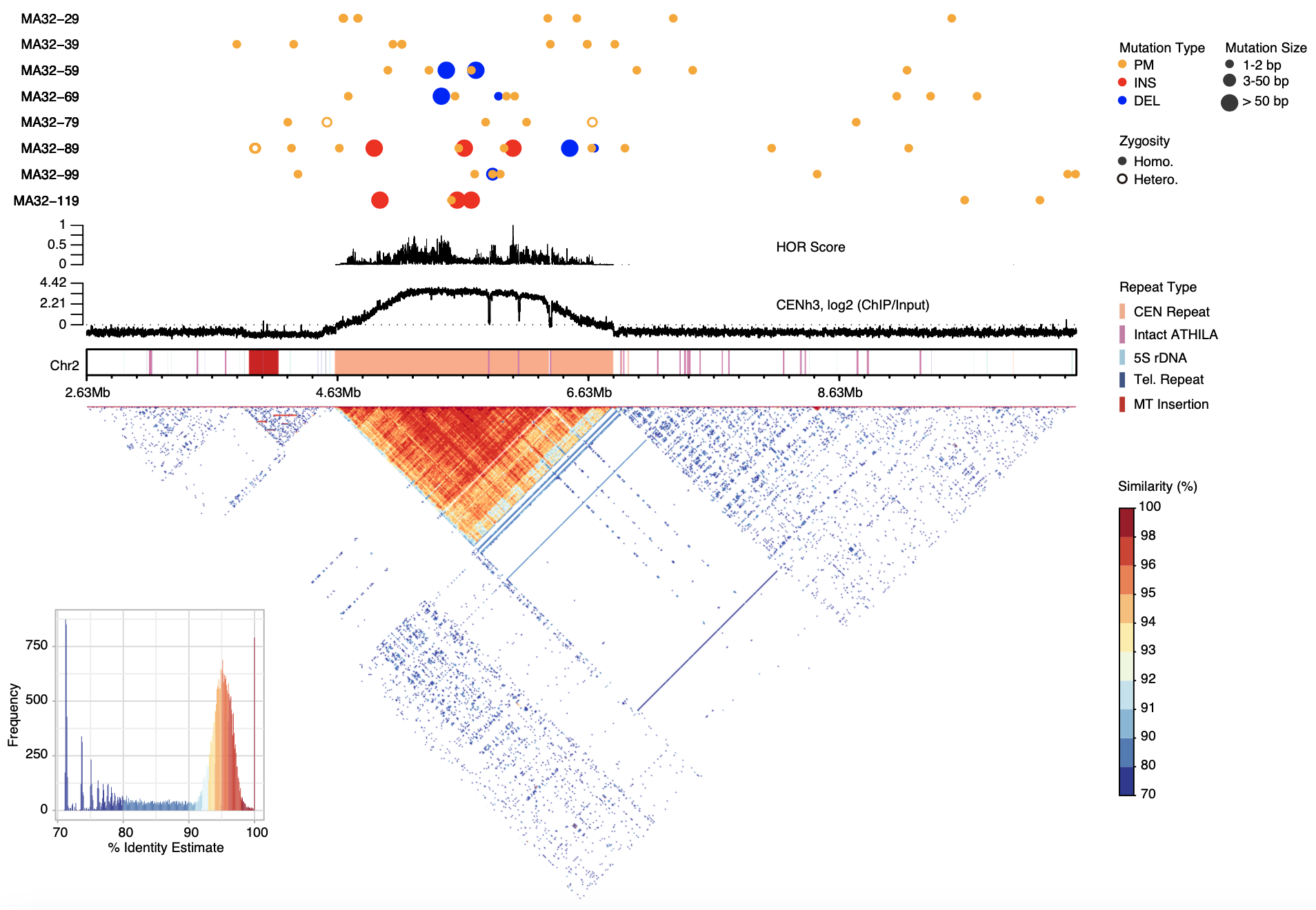

**Supplementary Fig. 13. Mutations within centromere 2.** Eight rows of circles display mutation patterns in the centromere of chromosome 2 in MA32 samples, colour-coded as point mutations (PMs; orange), insertions (red), and deletions (blue). Circle size reflects mutation length: small (1-2 bp), medium (3-50 bp), and large (>50 bp). Solid circles indicate homozygous mutations, while hollow circles indicate heterozygous ones. The two line plots below show higher-order repeat (HOR) scores and log2 CENH3 ChIP-seq enrichment, respectively. Rectangles highlight repetitive regions, and the heatmap below represents pairwise sequence identity between non-overlapping 10 kb regions, with a histogram summarizing identity values in the lower-left corner.

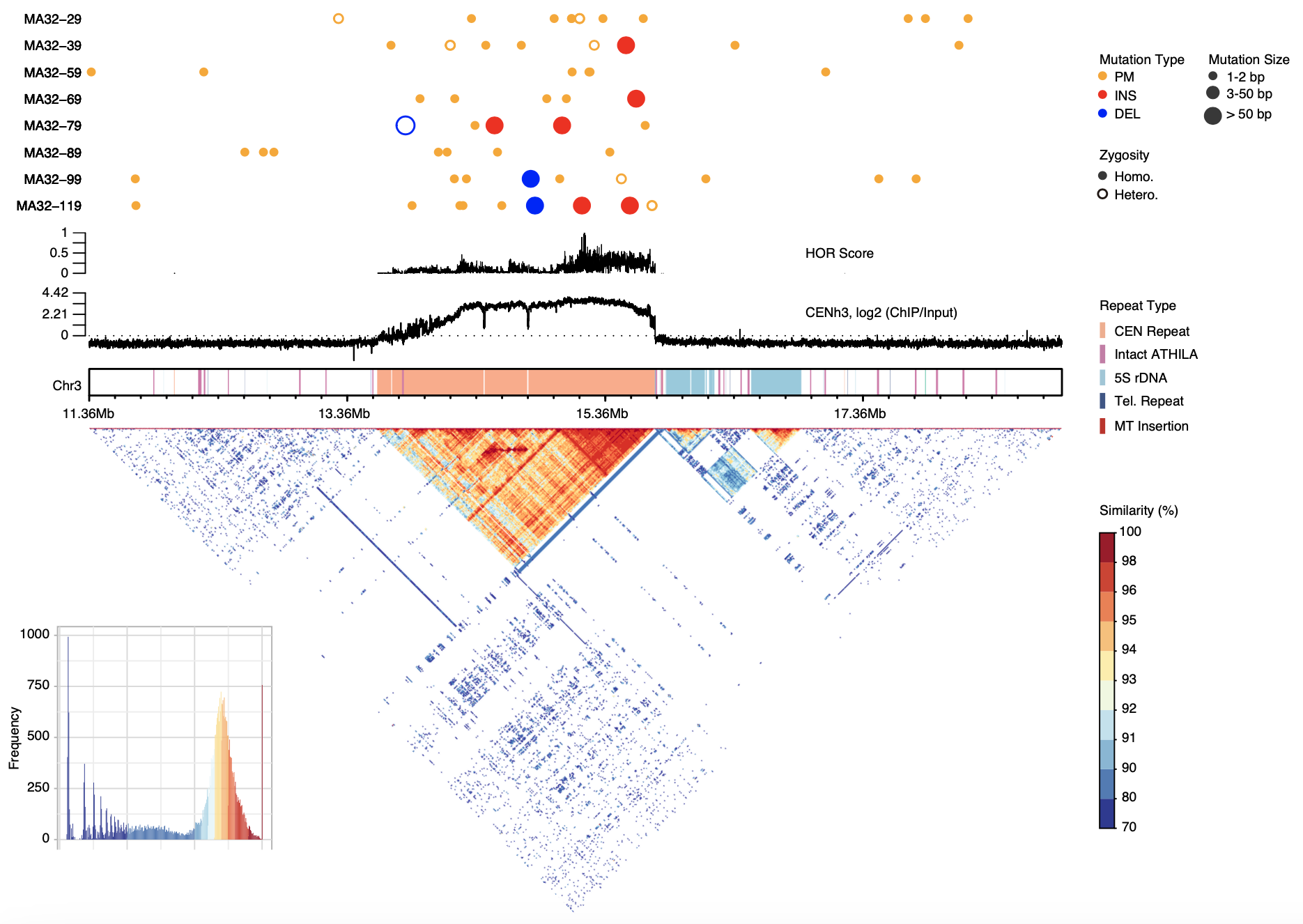

**Supplementary Fig. 14. Mutations within centromere 3.** Eight rows of circles display mutation patterns in the centromere of chromosome 3 in MA32 samples, colour-coded as point mutations (PMs; orange), insertions (red), and deletions (blue). Circle size reflects mutation length: small (1-2 bp), medium (3-50 bp), and large (>50 bp). Solid circles indicate homozygous mutations, while hollow circles indicate heterozygous ones. The two line plots below show higher-order repeat (HOR) scores and log2 CENH3 ChIP-seq enrichment, respectively. Rectangles highlight repetitive regions, and the heatmap below represents pairwise sequence identity between non-overlapping 10 kb regions, with a histogram summarizing identity values in the lower-left corner.

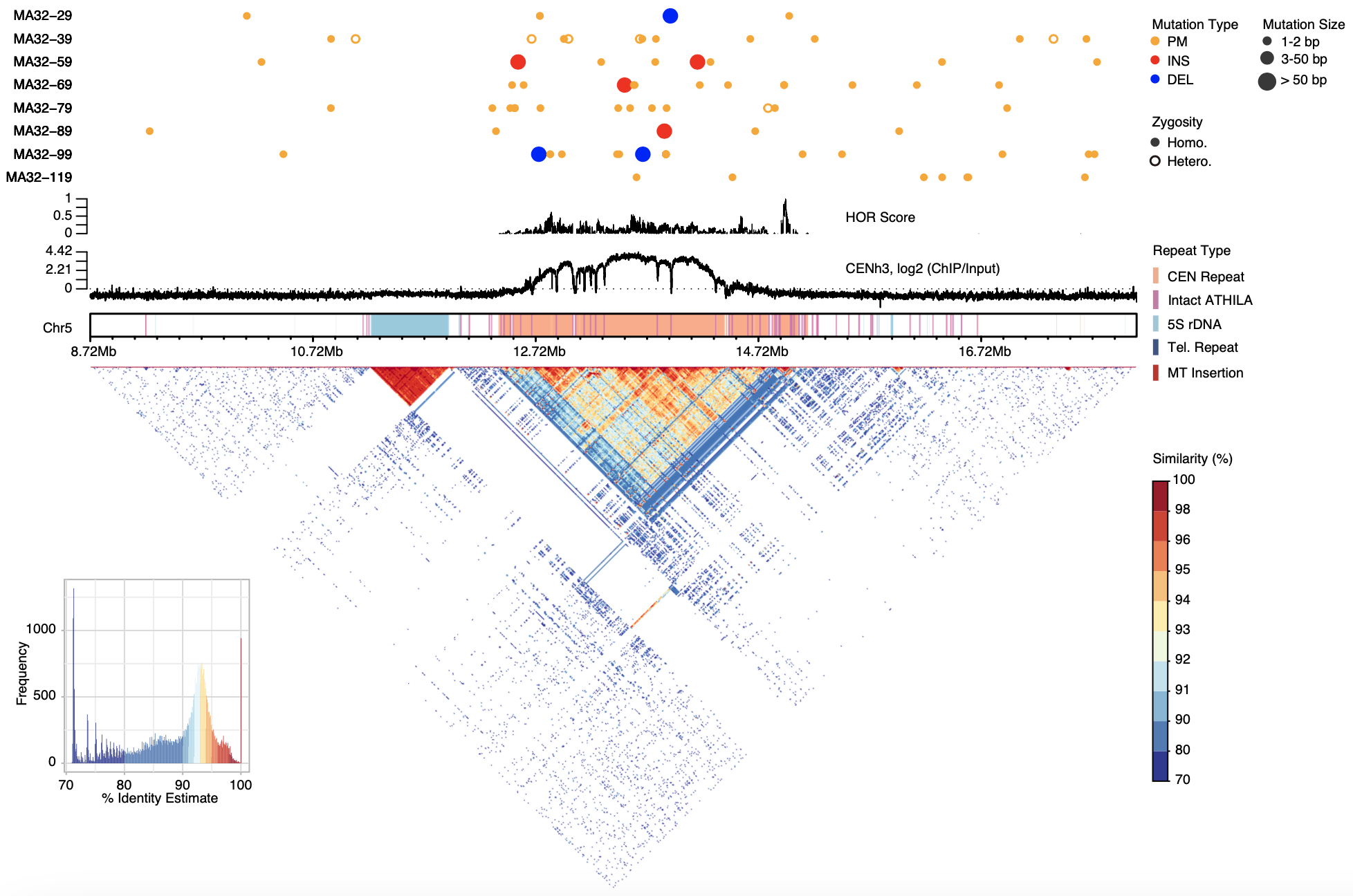

**Supplementary Fig. 15. Mutations within centromere 5.** Eight rows of circles display mutation patterns in the centromere of chromosome 5 in MA32 samples, colour-coded as point mutations (PMs; orange), insertions (red), and deletions (blue). Circle size reflects mutation length: small (1-2 bp), medium (3-50 bp), and large (>50 bp). Solid circles indicate homozygous mutations, while hollow circles indicate heterozygous ones. The two line plots below show higher-order repeat (HOR) scores and log2 CENH3 ChIP-seq enrichment, respectively. Rectangles highlight repetitive regions, and the heatmap below represents pairwise sequence identity between non-overlapping 10 kb regions, with a histogram summarizing identity values in the lower-left corner.

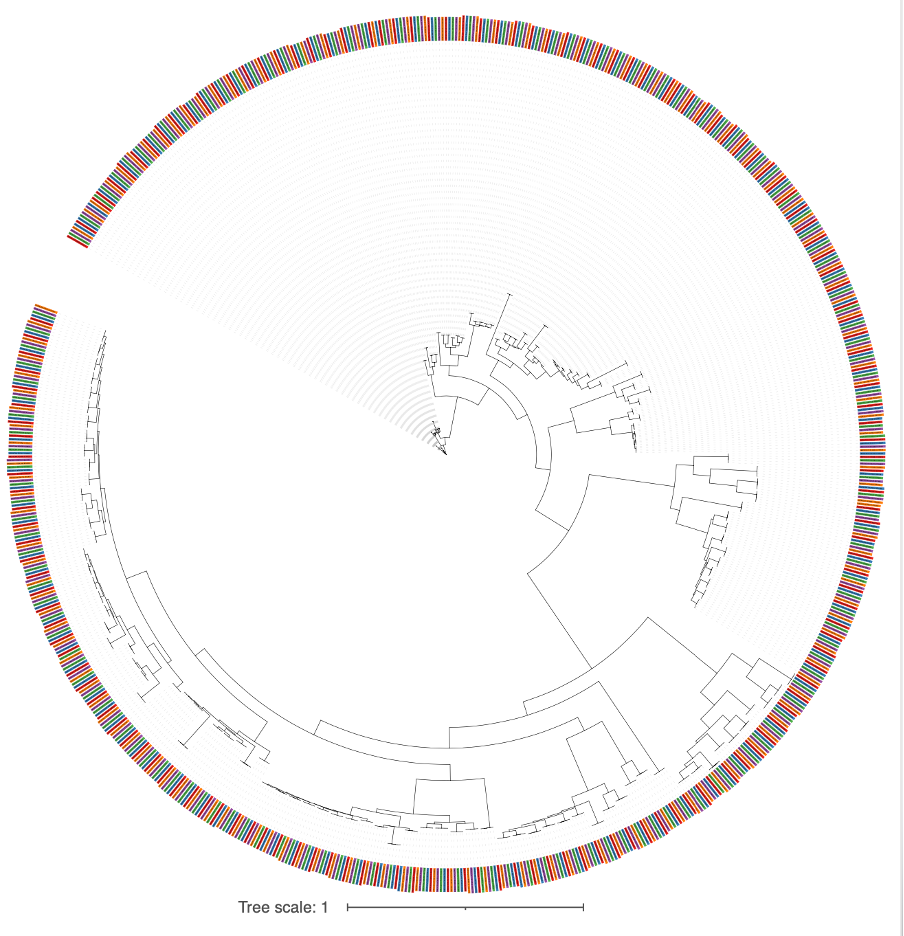

**Supplementary Fig. 16. Phylogenetic tree of 760 intact *ATHILA* elements from HPG1.** Each branch represents an individual element sequence, with labels coloured by sample. The robust clustering of five-member groups underscores the consistent identification and clear correspondence across samples.

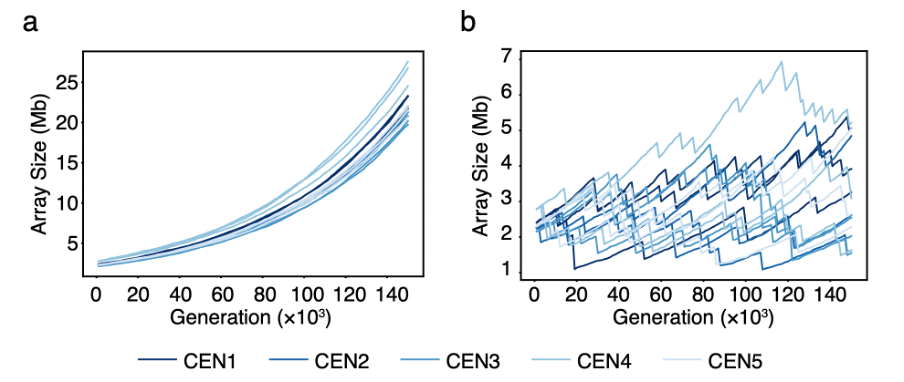

**Supplementary Fig. 17. Simulated size dynamics of five centromeric sequences.** **a.** Simulations based on mutation rates observed in the MA32 samples, incorporating increased numbers and lengths of insertion events. **b.** Simulations based on the mutation rates observed in MA32, with the addition of large deletions averaging 500 kb in size. Each centromeric sequence was simulated independently for 150,000 generations, with 20 (a) or 100 replicates (b) per sequence. Representative size trajectories from three simulations are shown.

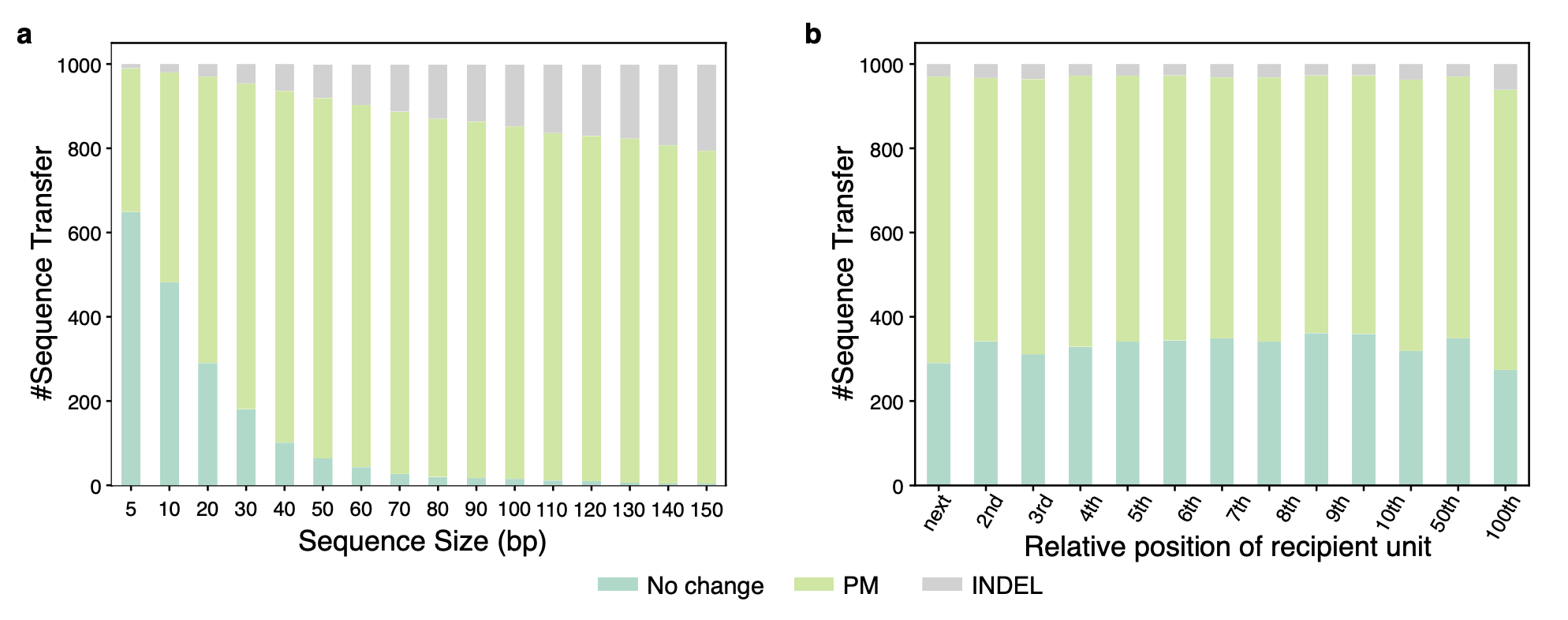

**Supplementary Fig. 18. Simulation of NAGC with different tract size (a) and distance (b).**

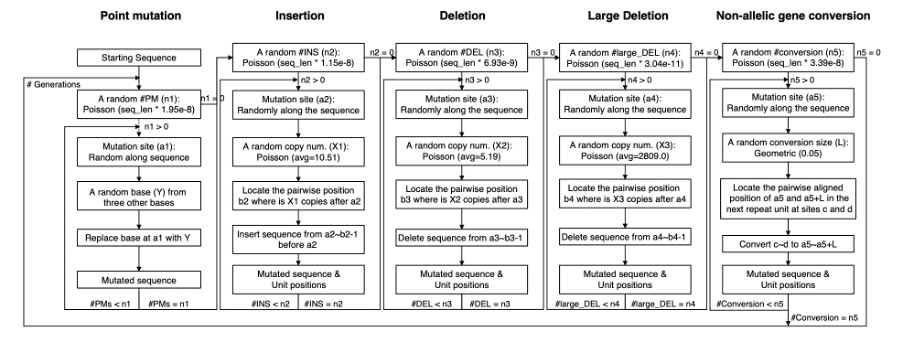

**Supplementary Fig. 19. Schematic overview of simulation pipeline.**
